# The Small RNA MicC is a Multifunctional Regulator of Extraintestinal Pathogenic *Escherichia coli* Fitness across Multiple Host Niches

**DOI:** 10.64898/2026.07.27.741068

**Authors:** O’Connor J. Matthews, Brittany A. Fleming, Alejandra A. Mendez, Kevin E. Nelson, Alan T. Stenquist, Sam Bonkowsky, Richard R. Kulesus, William J. Brazelton, Matthew G. Blango, Matthew A. Mulvey

## Abstract

Small non-coding RNAs (sRNA) modulate diverse bacterial functions ranging from carbon metabolism to virulence gene expression. Previous research showed that the sRNA chaperone Hfq is critical for the fitness of Extraintestinal Pathogenic *Escherichia coli* (ExPEC), a major cause of both bloodstream and urinary tract infections (UTI). Using the reference ExPEC strain UTI89, we created deletion mutants to probe the effects of seven conserved Hfq-dependent sRNAs (DsrA, RprA, OxyS, RyhB, MicF, MicC, Spf) on resistance to oxidative stress. All of the sRNA mutants grew normally in replete lysogeny broth, but the *spf* and *micC* mutants exhibited additive effects upon challenge with reactive oxygen species generated by methyl viologen. In a murine UTI model, the *spf* mutant resembled the wild-type strain, whereas UTI89Δ*micC* was unable to effectively colonize the bladder despite behaving like wild type within the kidneys. This correlated with a greatly reduced ability of the *micC* mutant to survive within bladder epithelial cells and paralleled UTI89Δ*micC* defects in gut colonization, virulence in a sepsis model, and complement resistance. Although MicC downregulated expression of its only known target, OmpC, aberrant modulation of this porin did not entirely account for the decreased stress resistance of UTI89Δ*micC*. Rather, RNA-Seq, sRNA target predictions, and *in vitro* phenotypic assays revealed that MicC can impact multiple pathways linked to niche establishment, including motility, chemotaxis, and various metabolic processes. These data are consistent with MicC serving as a multifunctional regulator of ExPEC stress responses and niche-specific fitness through OmpC-dependent and - independent mechanisms.

**IMPORTANCE:** Pathogenic strains of *Escherichia coli* are exceptionally common causes of diarrheal disease, urinary tract infection, sepsis, and meningitis. The ability of these pathogens to cause such a wide range of maladies is in part attributable to their ability to quickly adapt to and thrive within disparate and often hostile environments, including the gut, bladder, kidneys, and bloodstream. Adaptation to new environments requires rapid and precise changes in gene expression. To accomplish this feat, *E. coli* utilizes a suite of regulatory RNA called small RNA (sRNA). In this paper, we identified the sRNA MicC as a critical facilitator of *E. coli* fitness and virulence within diverse host environments via effects on the expression of multiple genes involved in bacterial motility, energy acquisition, and various other pathways. Delineating how sRNAs like MicC impact disease processes will aid the development of novel therapeutics to better combat *E. coli* infections.

## INTRODUCTION

Small non-coding RNA (sRNA) molecules help control the translation and activities of numerous proteins in both non-pathogenic and pathogenic bacteria (1–4). Base pairing with cognate mRNA targets enables sRNAs to act as molecular rheostats, tweaking mRNA translation levels up or, more often, down in response to varying environmental and nutritional cues by altering the accessibility of ribosome binding sites or by modulating mRNA stability. sRNAs can regulate diverse bacterial functions, including metabolism, iron utilization, the expression of outer membrane proteins (OMPs), stress and antibiotic resistance, quorum sensing, and virulence (1, 2, 5–8). Just over 100 different sRNA species have been confirmed in laboratory K-12 strains of *Escherichia coli* and, to a lesser extent, in their commensal and pathogenic counterparts (9). Pathogenic strains of *E. coli* typically have expanded genomes and are usually more stress resistant than laboratory K-12 strains, thus providing opportunities for the identification of new sRNA species and for expanding our functional understanding of even well-studied sRNAs (for examples, see (10–13)).

sRNA-mRNA interactions are oftentimes facilitated by the homohexameric RNA chaperone Hfq (14–16). Previously, our group reported that Hfq is critical for the fitness of Extraintestinal Pathogenic *E. coli* (ExPEC) (17). These bacteria are responsible for some of the most common and costly infections on the planet, including urinary tract infections (UTIs), bloodstream infections, and serious life-threatening sequelae like sepsis and meningitis (18, 19). Adding to the ExPEC threat is the rise and global dissemination of multidrug-resistant strains that greatly limit the effectiveness of many frontline antibiotics (19–21). Furthermore, the extensive use of antibiotics to treat common ExPEC-associated infections like UTIs creates selective pressures that likely promote the spread and amplification of resistance genes beyond the ExPEC lineage (22).

ExPEC strains are versatile, capable of adapting to a wide variety of rapidly changing and often hostile environmental conditions. These include the generation of reactive oxygen and nitrogen species by host cells, shifting nutrient limitations, antimicrobial peptides and other host defenses, and intense competition with resident microbes (23, 24). The latter is especially pertinent within the gastrointestinal tract, where ExPEC can co-exist with a multitude of other microbes, typically without eliciting any overt pathology (25). ExPEC strains can also persist within the genitourinary tract, where the pathogens can bind and invade mucosal epithelial cells (26, 27). Within the epithelial cells that comprise the bladder mucosa, ExPEC can replicate to form large, short-lived intracellular bacterial communities (IBCs) that eventually trigger the disruption and shedding of the host cells in which they reside. These processes facilitate the multiplication, local dissemination, and possibly the inter-host transmission of ExPEC (20, 27). IBCs may also enhance the establishment of small pockets of long-lived, and seemingly quiescent, intracellular reservoirs that can persist within the urinary tract even in the face of robust antibiotic treatments (27, 28). Growing evidence indicates that these intracellular reservoirs, along with gut-associated ExPEC communities, can seed the development of chronic and recurrent UTIs that vex many individuals throughout their lives (20, 25, 28, 29).

In our earlier work, deletion of *hfq* greatly compromised the ability of the ExPEC reference strain UTI89 to colonize and persist within the urinary tract, coincident with a reduced ability to form IBCs (17). Hfq also promoted UTI89 biofilm formation, motility, and resistance to various stressors, including reactive oxygen species (ROS). During a UTI, ROS are generated by bladder epithelial cells, macrophages, neutrophils, and other host cell types (30–35). ROS are also produced as by-products of bacterial metabolism, and these are elevated by the growth of some ExPEC strains within urine (36). Here, we set out to identify specific sRNA molecules that work in concert with Hfq to promote ROS resistance in ExPEC. To this end, we screened a panel of deletion mutants lacking seven previously documented Hfq-dependent sRNAs and identified Spot 42 (Spf) and MicC as important mediators of UTI89 ROS resistance. Subsequent assays revealed that MicC, but not Spf, was important for UTI89 persistence within the bladder, and specifically within bladder epithelial cells. In addition, we report that MicC is critical for UTI89 persistence within the gut and absolutely required for ExPEC virulence in a murine model of sepsis, while Spf was entirely dispensable in these assays. Currently, the only validated target of MicC in *E. coli* is the transcript for the porin OmpC (37). Here, using RNA-Seq and other approaches, we show that changing MicC levels in UTI89 can alter multiple processes beyond OmpC expression, including varied metabolic pathways and motility. These results help explain the broad reaching effects of MicC on UIT89 stress resistance and virulence phenotypes, providing direction for future research aimed at delineating specific MicC binding partners within ExPEC while also highlighting the potential value of sRNAs like MicC as therapeutic targets.

## RESULTS

### Spot 42 and MicC independently enhance ExPEC resistance to oxidative stress

To assess how Hfq might promote ExPEC resistance to ROS, we examined seven previously described Hfq-dependent sRNAs that are highly conserved among all *E. coli* strains and have one or more known targets directly or indirectly modulating oxidative stress resistance (**Table 1**). Two of the sRNAs, DsrA and RprA, promote expression of the stationary-phase sigma factor RpoS (σ^S^) (38, 39), while another (OxyS) inhibits RpoS expression and can help protect lab K-12 *E. coli* strains against ROS (40–43). MicC and MicF offer two examples of sRNA molecules that can regulate OMPs, repressing the synthesis of OmpC and OmpF, respectively (37, 44). Both MicC and MicF are induced upon exposure to superoxide (37). Reduction of OMPs may decrease the ability of ROS to breach the bacterial cell wall. The sRNA RyhB regulates the biosynthesis of siderophores and Fe-S clusters (which are highly sensitive to oxidative stress) and is repressed by the iron-sensing regulator Fur (12, 45, 46). The prototypical sRNA Spot 42 (Spf) is known to modulate basal metabolic processes and may indirectly alter levels of metabolically produced ROS (47, 48). Spf, which is an especially abundant sRNA, can also affect the activities of other sRNA molecules by competing for Hfq binding sites (49). In addition to the Spf sRNA, the *spf* locus also encodes a small 15-amino acid peptide that can bind to and inhibit the functionality of the cAMP receptor protein CRP (50), which is critical for ExPEC stress resistance and survival within the urinary tract (51). Each of these sRNAs was deleted individually in UTI89 using lambda-Red-mediated homologous recombination (52).

**Table 1.** sRNA molecules assayed for effects on ExPEC growth under oxidative stress.

| Name | Length (UTI89) | % identity of UTI89 sequence with K-12 *E. coli* (MG1655) | Known Targets in *E. coli* * | Reference |
| --- | --- | --- | --- | --- |
| DsrA | 86 | 100 | *mreB*, *hns*, *rbsD*, ***rpoS*** | (38, 146, 147) |
| MicC | 108 | 97 | *ompC* | (37) |
| MicF | 92 | 99 | *ompF*, *cpxR*, *lrp*, *obgE*, ***oppA***, *phoE*, *seqA* | (148-152) |
| OxyS | 109 | 100 | *fhIA*, *flhDC*, *iscR*, *mepS*, *nusG*, *rpoS* | (42, 43, 153-156) |
| RprA | 104 | 100 | *csgD*, *fepA*, *hdeD*, ***rpoS***, *ydaM* | (39, 157-159) |
| RyhB | 89 | 100 | *acnA*, *acnB*, *bfr*, ***cirA***, *cydA*, *cysE*, *dhaKLM*, *dmsA*, *dmsD*, *erpA*, *fepB*, *fumAC*, *fumB*, *gltB*, *grxD*, *hemB*, *hemH*, *hscBA-fdx-iscX*, *iscRSUA*, *katG*, *leuZ*, *msrB*, *napF*, *nirB*, *nrfA*, *nuo* genes, *sdhCDAB*, ***shiA***, *sodB*, *sufABCDSE*, *yegD*, *ynfEFGGH-ykgJ* | (45, 46, 159-161) |
| Spot 42 (Spf) | 108 | 100 | *araF*, *ascF*, *atoD*, *caiA*, *fucI*, *fucP*, *galK*, *gdhA*, *glpF*, *gtlA*, *icd*, *nanC*, *nanT*, *paaK*, *puuE*, *sdhC*, *sepL*, *srlA*, *sthA*, *sucC*, *xylF* | (47, 48, 148, 162-165) |

\*Genes in bold indicate targets that are activated by the indicated sRNA; all other targets are repressed.

Sensitivity to oxidative stress was tested using methyl viologen (MV), which generates superoxide radicals when added to lysogeny broth (LB) cultures (53). All the sRNA deletion strains grew similarly to the wild-type UTI89 control strain in LB alone (**Fig. 1A**). In contrast, the *micC* and *spf* knockouts had clear and highly reproducible growth defects in the presence of 1 mM MV (**Fig. 1B**). A double knockout lacking both *spf* and *micC* rendered UTI89 more sensitive to MV than either single deletion alone, suggesting that these two sRNAs promote oxidative stress resistance via effects on independent pathways (**Fig. 1C**). In line with this conclusion, *micC* and *spf* show negative epistasis as assessed by use of a multiplicative Bliss model (54). Of note, OD_600_ readings in our growth assays correlated well with viable bacterial numbers, as determined by quantifying colony forming units (CFU)/ml via dilution plating (**Fig. 1D–G**). Leaky expression of plasmid-borne *spf* downstream of a P*tac* promoter rescued growth of the *spf* knockout in the presence of MV (**Fig. 1H**). The *micC* knockout could be similarly complemented dependent upon the level of induction by IPTG; low levels of induction rescued growth while higher levels of induction were deleterious (**Fig. 1I**). Collectively, these results indicate that Spf and MicC promote ExPEC resistance to oxidative stress using at least partially distinct mechanisms.

**Figure 1.**
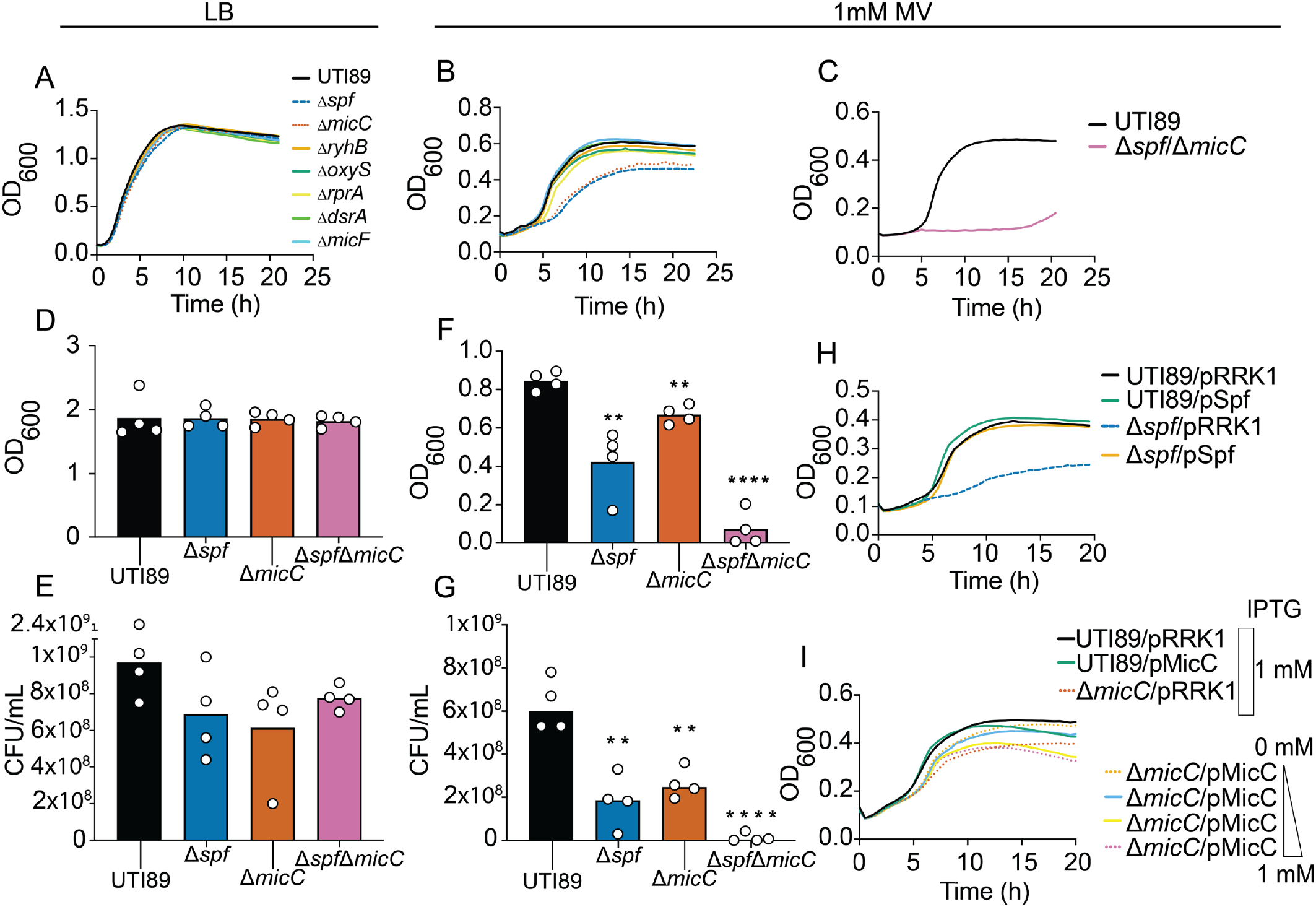
UTI89Δ*micC and* UTI89Δ*spf* have increased sensitivity to ROS. **(A-C)** Curves generated using a Bioscreen C instrument show growth of wild-type UTI89 and isogenic sRNA knockouts in **(A)** LB or **(B-C)** LB + 1 mM MV. **(D-E)** OD_600_ of UTI89, UTI89Δ*spf*, UTI89Δ*micC*, and UTI89Δ*spfΔmicC* cultures after 16 h of growth in 5 ml **(D)** LB or **(E)** LB + 1 mM MV. **(F and G)** Titers of viable bacteria (CFU/ml) recovered from (**F**) LB and (**G**) LB + 1 mM MV cultures after 16 h (same cultures as in **(D)** and **(E)**, respectively). **(H)** Curves show growth of UTI89 and UTI89Δ*spf* carrying the empty vector pRRK1 or pSpf in LB containing 1 mM MV and 1 mM IPTG. **(I)** Curves depict growth of UTI89 and UTI89Δ*micC* carrying the empty vector pRRK1 or pMicC in LB plus 1 mM MV. IPTG was added at a concentration of 1 mM or in 10-fold increments from 1 mM down to 0 mM, as indicated in the legend. Each growth curve **(A-C, H, I)** represents the means of quadruplicate samples and are representative of experiments performed three or more times. Error bars for the growth curves were minor and omitted for clarity. Bar graphs **(D-G)** represent the mean of 4 independent experiments (indicated by dots). *P* values were determined by Student’s *t* tests; \*\**P* < 0.01, ****P < 0.0001, versus the wild-type control.

### MicC promotes ExPEC colonization of the bladder

The ability of ExPEC to resist oxidative stress is important for bacterial survival during a UTI (36, 51, 55–60). As MicC and Spf promoted ExPEC resistance to ROS in our *in vitro* assays (**Fig. 1**), we next asked if these sRNAs impact ExPEC fitness within the urinary tract. For this, we used a well-established murine UTI model in which adult female CBA/J mice were inoculated via transurethral catheterization with ∼10^7^ CFU of wild-type UTI89, UTI89Δ*spf*, or UTI89Δ*micC* (28). Comparable numbers of wild-type UTI89 and UTI89Δ*spf* were recovered from the bladders and kidneys at 3- and 5-days post-inoculation (dpi) (**Fig. 2A-D**). In contrast, at these time points UTI89Δ*micC* had a clear defect within the bladder but was recovered at near wild-type levels from the kidneys. Similar results were observed in C3H/HeJ mice at 3 dpi (**Fig. 2E-F**). Due to a mutation in the gene for the pathogen recognition receptor Toll-like Receptor 4 (TLR4), and possibly defects in other factors, C3H/HeJ mice have reduced inflammatory responses, and neutrophils recovered from these animals are impaired in their ability to generate ROS (61–64). In total, our findings indicate that MicC is an important fitness determinant for ExPEC within the bladder, even in the presence of attenuated host inflammatory responses.

**Figure 2.**
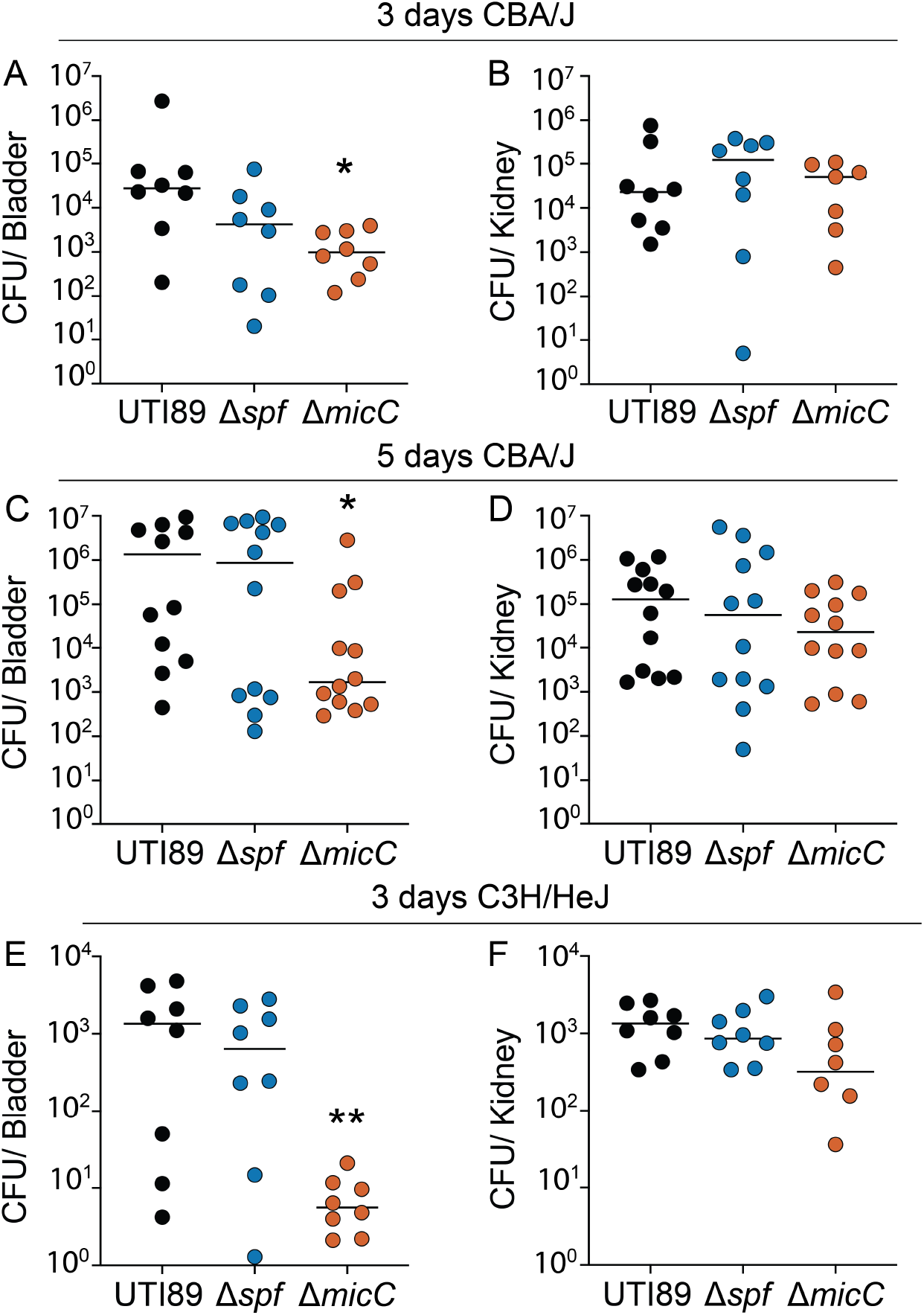
MicC promotes ExPEC fitness within the murine bladder. The bladders of adult female **(A-D)** CBA/J mice or **(E-F)** C3H/HeJ were inoculated with ∼10^7^ CFU of UTI89, UTI89Δ*spf*, or UTI89Δ*micC* via transurethral catheterization. Mice were sacrificed 3 d **(A-B, E-F)** or 5 d **(C-D)** post-inoculation and bacterial titers within the bladders and kidneys were determined by plating tissue homogenates. Bars in all graphs denote median values. \**P<*0.05; \*\**P*<0.01 by Mann Whitney U tests. *n* ≥ 7 mice per group.

### MicC is important for ExPEC persistence within bladder epithelial cells

Considering the diminished capacity of UTI89Δ*micC* to colonize the bladder, we next assessed the ability of the *micC* mutant to bind, invade, and survive within bladder epithelial cells (BECs) relative to wild-type UTI89 and UTI89Δ*spf*. In experiments using the 5637 line of human BECs, UTI89Δ*micC* was significantly impaired in its ability to associate with the host cells in comparison with UTI89 and UTI89Δ*spf* (**Fig. 3A**). In parallel, gentamicin protection assays indicated that the Δ*micC* mutant was unexpectedly much better than the wild-type strain at invading the BECs (**Fig. 3B**), but then failed to persist intracellularly (**Fig. 3C**). As in the murine UTI model, no significant differences were observed between wild-type UTI89 and UTI89Δ*spf* in any of these cell culture-based experiments. The markedly reduced ability of UTI89Δ*micC* to persist within BECs may help explain the bladder-specific defects observed with this mutant in the murine UTI model.

**Figure 3.**
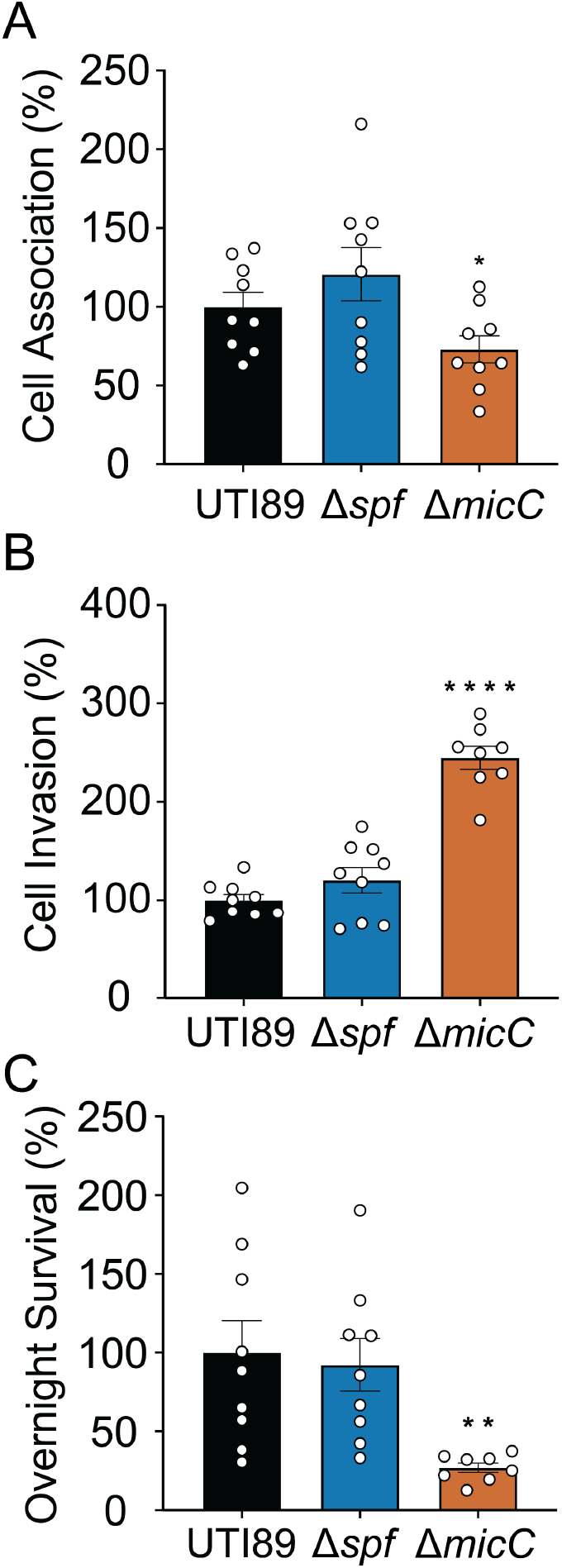
MicC impacts the ability of ExPEC to associate with, invade, and survive within BECs. Human BECs were infected with UTI89, UTI89Δ*spf*, and UTI89Δ*micC* for 1.5 h followed by additional 1-h and 17.5-h incubations in the medium containing gentamicin (100 μg/ml). **(A)** Graph shows relative levels of host cell-associated bacteria prior to the addition of gentamicin, normalized to the total numbers of bacteria present at the 1.5-h time point. Graphs in **(B-C)** indicate levels of viable, intracellular bacteria recovered at **(B)** 1 h and **(C)** 17.5 h after the addition of gentamicin. Data are expressed relative to wild-type UTI89, with bars indicating median values (± SEM) from 3 independent experiments carried out in triplicate. Invasion rates were calculated relative to the numbers of host cell-associated bacteria present prior to the addition of gentamicin, while overnight survival data were normalized to invasion rates. *P* values were determined by Welch’s *t* tests; \**P* < 0.05, \*\**P* < 0.01, \*\*\*\**P* < 0.0001.

### MicC provides a competitive advantage within the gut

To determine the effects of MicC and Spf on the ability of UTI89 to colonize the gut, which serves as a major reservoir for ExPEC that can seed UTIs and other extraintestinal infections (25, 65), adult specific-pathogen-free (SPF) BALB/c mice were inoculated via oral gavage with 10^9^ CFU each of UTI89, UTI89Δ*spf*, or UTI89Δ*micC*. Colonization levels were then tracked by collecting, homogenizing, and plating feces on selective media at various time points over the course of 3 weeks. For these assays, UTI89 and the *micC* and *spf* deletion mutants were engineered to express either kanamycin (Kan^R^) or chloramphenicol (Cam^R^) resistance cassettes so that they could be easily distinguished (65–67). In this model, the numbers of viable ExPEC recovered from the feces correlate well with ExPEC titers within the large intestines (65). All 3 strains effectively colonized the murine intestinal tract and persisted at fairly similar levels for at least 3 weeks (**Fig. 4A**). To further test the fitness of these strains within the gut, we employed competitive assays in which the wild-type strain was mixed 1:1 with either UTI89Δ*spf* or UTI89Δ*micC* prior to delivery into mice via oral gavage. In these assays, the wild-type strain and UTI89Δ*spf* co-existed at similar levels until 14 dpi, at which point the Δ*spf* mutant was recovered at levels that were about 25-fold higher than the wild-type strain, reflecting a median competitive index of 1.55 (**Fig. 4B**). In contrast, starting at 7 dpi UTI89Δ*micC* was clearly outcompeted by wild-type UTI89 (**Fig. 4C**). By day 14, there was about a 120-fold reduction in the relative levels of UTI89Δ*micC* recovered in the feces, reflecting a median competitive index of -2.08. These results indicate that loss of *micC* greatly impairs the competitive fitness of ExPEC within the gut, while the deletion of *spf* provides a perceptible advantage.

**Figure 4.**
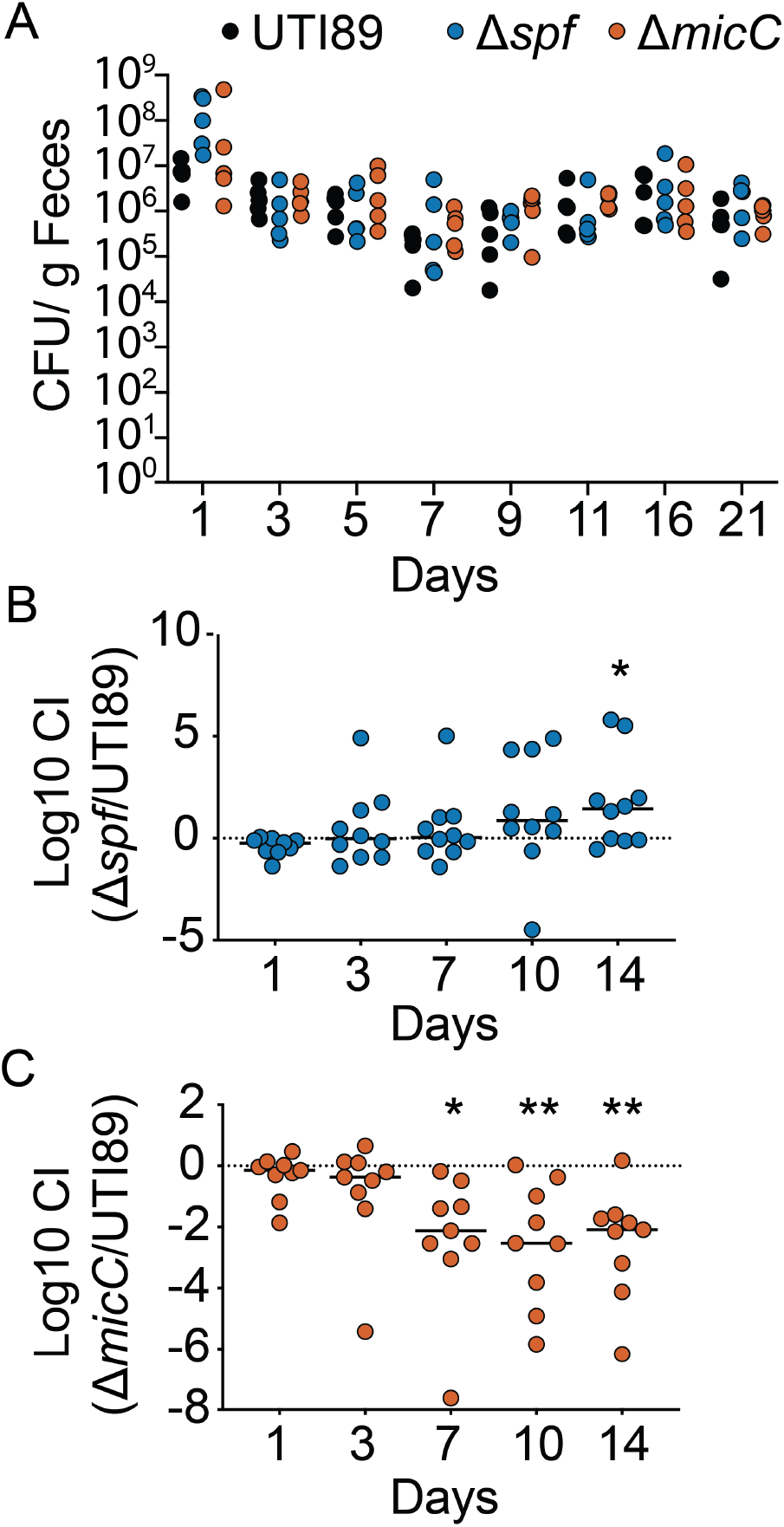
MicC promotes gut colonization by ExPEC. **(A)** Graph shows results from noncompetitive assays in which Balb/C mice were inoculated with 10^9^ CFU of UTI89, UTI89Δ*spf*, or UTI89Δ*micC* by oral gavage, and fecal titers were determined at the indicated time points (n = 5 mice). For competitive assays, mice were inoculated with ∼10^9^ CFU of a 1:1 mixture of **(B)** wild-type UTI89 and UTI89Δ*spf* or **(C)** UTI89 and UTI89Δ*micC.* Fecal titers of each strain were determined at the indicated time points by plating on selective medium. \**P*<0.05; \*\**P*<0.01 by one sample *t*-tests. Bars denote median values; n = 9-10 mice from two independent assays.

### MicC enhances serum resistance and is critical for ExPEC virulence within the bloodstream

ExPEC strains are the principal cause of bloodstream infections and a leading cause of sepsis (68–70). The entry of ExPEC into the bloodstream appears to occur predominantly via either translocation across the intestinal mucosa or by dissemination from the urinary tract (70–73). Once within the bloodstream, ExPEC faces nutrient limitations and a torrent of host defenses including patrolling phagocytes, antibodies, and the complement system. Only a subset of ExPEC isolates appear capable of surviving and causing severe disease following entry into the bloodstream (73–76). To test if the sRNAs Spf and MicC are important for ExPEC virulence within the bloodstream, adult outbred female Swiss-Webster mice were inoculated intraperitoneally with 10^6^ CFU of UTI89, UTI89Δ*spf*, or UTI89Δ*micC* and then monitored for signs of morbidity. With this route of inoculation, ExPEC rapidly enter the bloodstream and disseminate (67, 77). In our experiments, all mice infected with UTI89 and UTI89Δ*spf* perished by 24 h post-inoculation (hpi) (**Fig. 5A**). In sharp contrast, all mice infected with UTI89Δ*micC* survived beyond 48 hpi, indicating an important function for MicC in promoting ExPEC virulence during bloodstream infections.

**Figure 5.**
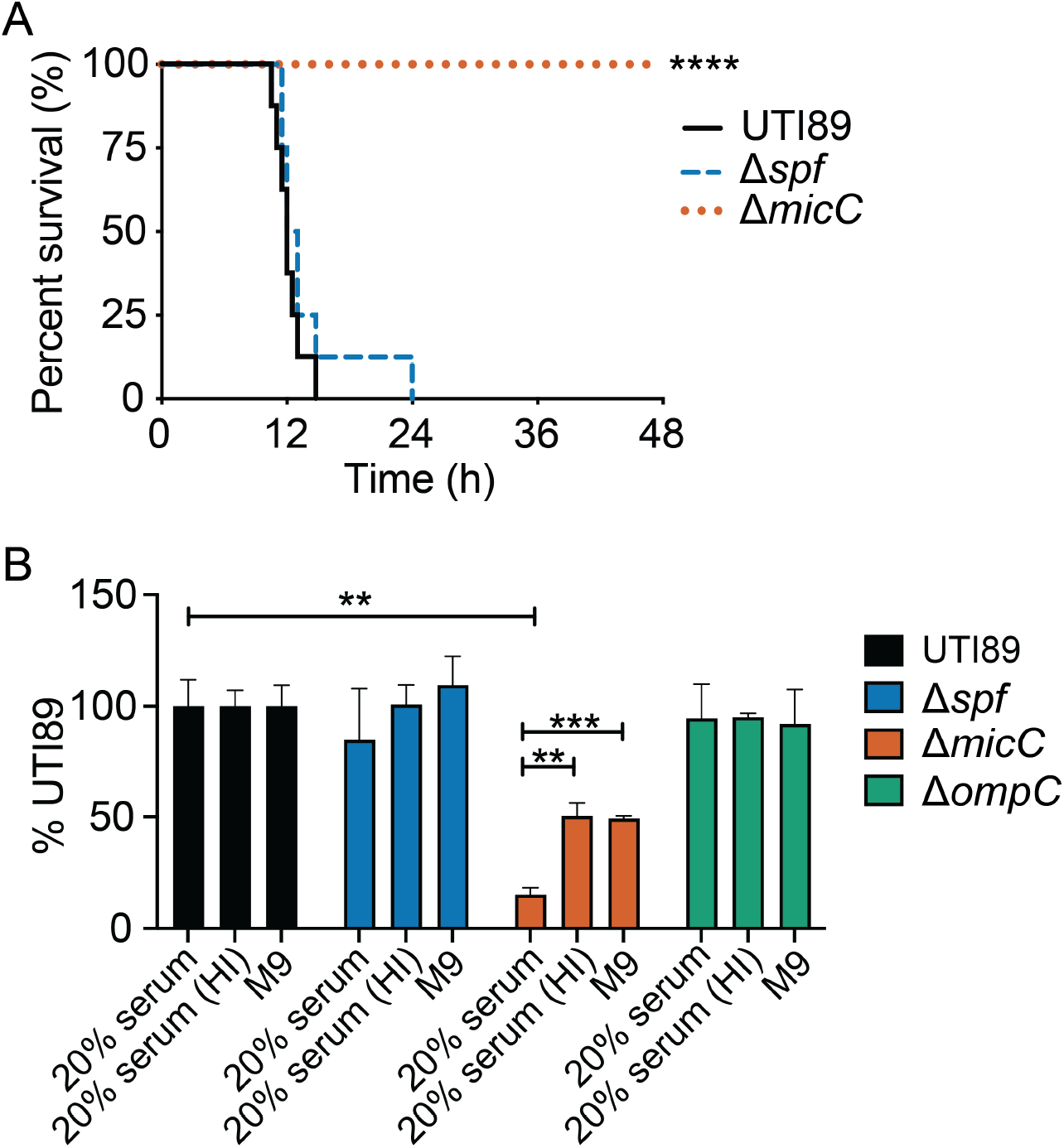
MicC is important for ExPEC virulence in the bloodstream and serum resistance. **(A)** Kaplan Meier survival curves of Swiss Webster mice inoculated with 10^6^ CFU intraperitoneally. \*\*\*\**P*<0.0001 by Log-rank Mantel Cox test. *n* = 8-9 mice per group. **(B)** Titers of UTI89, UTI89Δ*spf,* UTI89Δ*micC*, UTI89Δ*ompC* after 2.5 h of growth in M9 medium supplemented with 20% pooled human serum or 20% heat-inactivated (HI) pooled human serum, normalized to input titers and control. Bar graphs represent the mean of three independent experiments, ± SEM. *P* values were determined by Student’s *t* tests; \*\**P*<0.005 and *\*\*\*P*<0.0005, versus the wild-type control.

As UTI89Δ*micC* was avirulent in our sepsis model, we wanted to determine the sensitivity of this mutant to complement-mediated killing. The complement system, which localizes within the serum fraction of blood, is composed of multiple heat-labile factors that can mediate bacterial opsonization and the formation of membrane attack complexes that can trigger bacterial lysis (78). To assess the relative sensitivities of wild-type UTI89, UTI89Δ*spf*, and UTI89Δ*micC* to complement-mediated killing, we performed serum resistance assays in which the bacteria were incubated for 2.5 h in M9 minimal media supplemented with 20% pooled human serum. In these assays, UTI89 and UTI89Δ*spf* grew similarly in our standard M9 media (which contained both glucose and casein amino acids as carbon sources), and both strains displayed comparable levels of serum resistance (**Fig. 5B**). Assessing the serum resistance of UTI89Δ*micC* was complicated because this mutant grew less robustly than UTI89 and UTI89Δ*spf* in the M9 media that was used for these assays. This reflects metabolic defects in UTI89Δ*micC* that are considered in more detail later. Nonetheless, in our serum resistance assays UTI89Δ*micC* titers were clearly reduced in the presence of active serum (**Fig. 5B**). Of note, the use of heat-inactivated (HI) serum, which lacks functional complement, had no effect on any of the tested strains.

The only bonified target of MicC in *E. coli* is the outer membrane protein OmpC (37). In a previous study, it was reported that an *E. coli* clinical isolate (strain 2837-2/05) lacking OmpC was substantially more resistant to complement-mediated killing in comparison to its wild-type counterpart (79). In light of this information and considering the regulatory link between MicC and OmpC (37), we tested the effect of OmpC on UTI89 survival in our serum-resistance assays. In these experiments, the deletion of *ompC* had no discernable effect on UTI89 survival (**Fig. 5B**). Our results may contrast with the previously published findings with *E. coli* 2837-2/05 due to differences in the strains or methods used (e.g the use of 20% serum here versus 60% in the earlier study) (79). In particular, bacterial killing in the previous study was mediated by the classical complement pathway requiring anti-OmpC antibodies, which may have been limiting in our assays.

### Varying OmpC levels alter ExPEC sensitivity to ROS

The lack of any phenotype with the *ompC* mutant in our serum resistance experiments prompted us to consider how OmpC and MicC might be phenotypically linked in other assays with UTI89. MicC inhibits the translation of OmpC by binding to sequences within the 5’ untranslated region of *ompC* transcripts (37). By analyzing outer membrane fractions via SDS-PAGE, we found that overexpression of MicC led to the clear downregulation of OmpC in UTI89 without affecting the expression of the OmpA and OmpF porins (**Fig. 6A**). These results are in agreement with observations made in the lab K-12 *E. coli* strain BW25113 that initially implicated MicC as a negative regulator of OmpC expression (37). In this earlier study, OmpC levels were modestly elevated in BW25113 in the absence of MicC, but we did not see this effect consistently in our experiments with UTI89 (**Fig. 6B**), possibly due to input from other OmpC regulators (80, 81).

**Figure 6.**
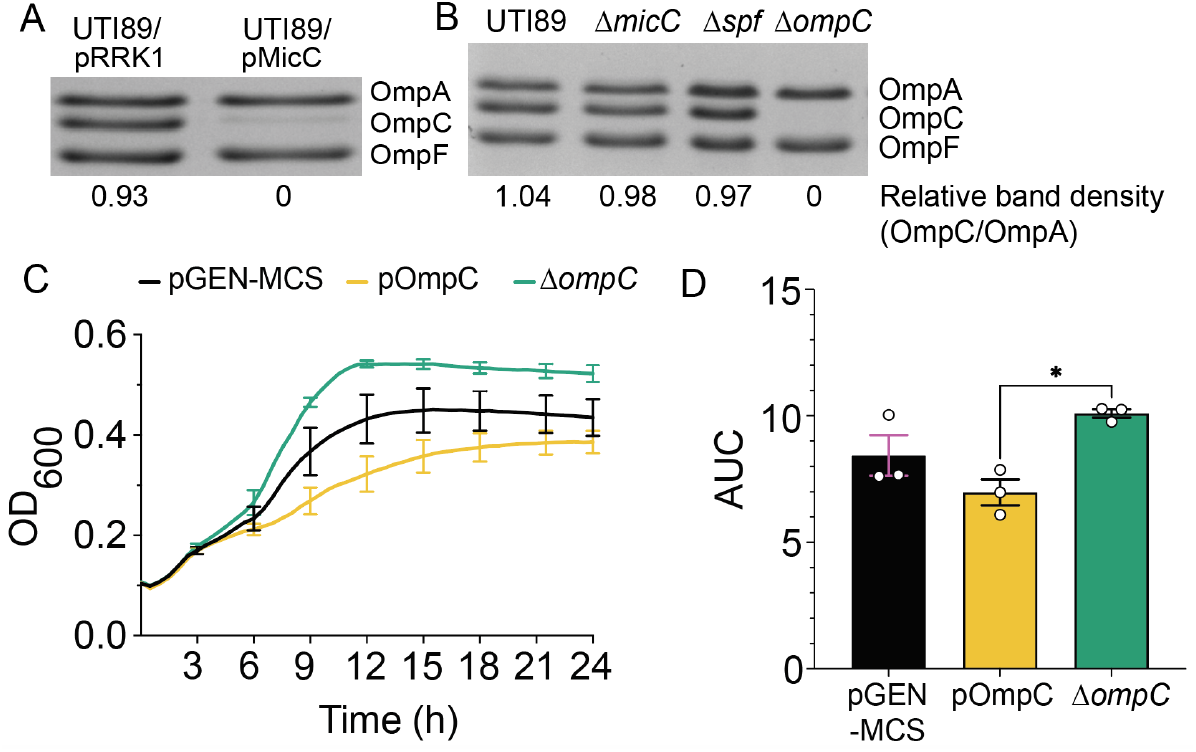
OmpC effects on MV sensitivity of ExPEC. **(A)** Coomassie blue-stained gel shows levels of the major OMPs present in outer membrane fractions isolated from UTI89 carrying the empty vector pRRK1 or pMicC, following growth to mid-log in LB + 1 mM IPTG (to induce expression of MicC). **(B)** Gel shows levels of major OMPs recovered from UTI89, UTI89Δ*micC*, UTI89Δ*spf*, and UTI89Δ*ompC* following after reaching OD_600_ of 0.6. Relative levels of OmpC, as determined by densitometry measurements, are indicated. **(C)** Curves indicate growth of UTI89/pGEN-MCS (control empty vector), UTI89/pOmpC, and UTI89Δ*ompC* in LB + 1 mM MV. **(D)** Bars show mean Area Under the Curve (AUC) values the growth data presented in (C). Each curve in (C) is an average of 3 independent biological replicates carried out with 3 or more samples. Error bars indicate SEM; \**P* < 0.05 as determined using Welch’s *t* tests.

We next asked if changing OmpC levels could recapitulate the defects observed with UTI89Δ*micC* grown in the presence of ROS generated by the addition of MV (see **Fig. 1**). The deletion of *ompC* markedly enhanced the growth of UTI89 in LB containing MV, while the overexpression of OmpC had an inhibitory effect (**Fig. 6C and D**). The latter phenotype mirrors that seen with growth of UTI89Δ*micC* in the presence of MV (see **Fig. 1**). However, since the deletion of *micC* had little effect on OmpC levels in UTI89 (**Fig. 6B**), it is likely that the increased sensitivity of UTI89Δ*micC* to MV is not entirely attributable to the mis-regulation of OmpC expression.

### Perturbation of MicC levels causes widespread transcriptional changes

To examine how changing MicC levels might impact ExPEC beyond effects on OmpC, we used RNA-Seq to analyze the transcriptomes of UTI89/pRRK1 relative to UTI89Δ*micC*/pRRK1 and UTI89Δ*micC*/pMicC. In these assays, pRRK1 was included as an empty vector control for the IPTG-inducible MicC expression construct pMicC. Bacterial cultures were grown to mid-log phase prior to the addition of IPTG for 1.25 h and subsequent processing for RNA-Seq. Relative to UTI89/pRRK1 (which served as the wild-type, WT, control for these experiments), 1,475 genes were differentially expressed (*P adjusted* ≤ 0.05) in the knockout (KO) mutant UTI89Δ*micC*/pRRK1 (**Table S1**). Among these, 171 were expressed greater than two-fold in the *micC* mutant, while 340 were downregulated more than two-fold in relation to UTI89/pRRK1 (**Fig. 7A**). In the MicC overexpression strain (OE, UTI89Δ*micC*/pMicC), the levels of 247 transcripts were significantly altered relative to the WT strain, with 176 varying by at least two-fold (**Fig. 7B**). 104 transcripts were differentially expressed in both the KO and OE strains relative to UTI89/pRRK1, and among these 49 varied by at least two-fold between the KO and OE strains (**Table 2**, **Table S1**, sheets labeled ‘Shared Hits (all)’ and ‘Shared Hits (abs log_2_ diff ≥ 1)’). This latter group of genes include the only validated MicC target in *E. coli*, *ompC*, which was down about 4.4-fold in the OE strain and 1.5-fold in the *micC* KO.

**Figure 7.**
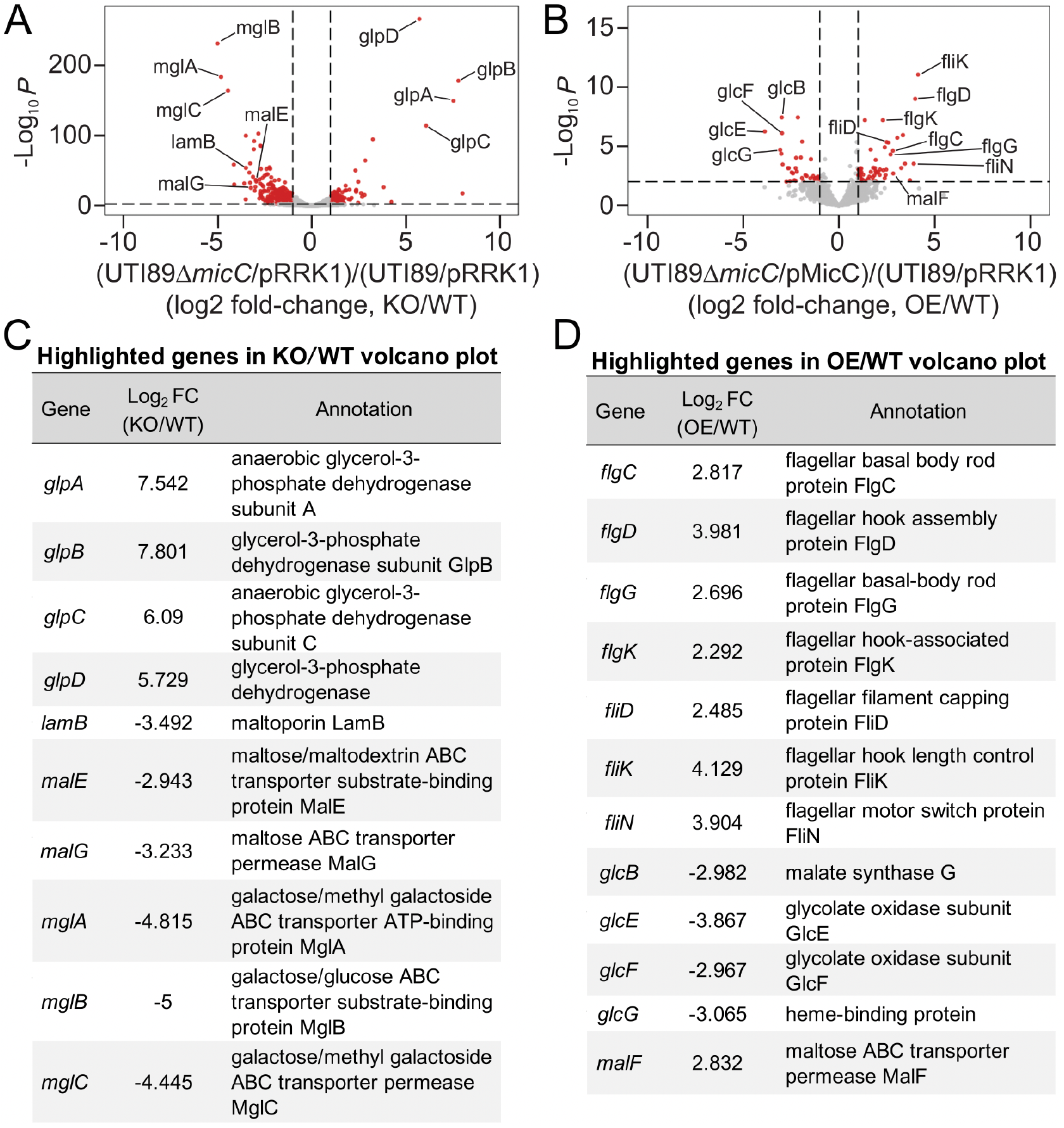
MicC has widespread effects on ExPEC gene expression. (A and. **B)** Volcano plots show relative transcript levels (Log_2_-fold change) versus *P* values (-Log_10_) in UTI89Δ*micC/*pRRK1 relative to UTI89/pRRK1 (KO/WT), or in UTI89Δ*micC*/pMicC relative to UTI89/pRRK1 (OE/WT), as assessed by RNA-seq. For these assays, overnight cultures were back-diluted 1:100 in LB and grown for 2 h shaking at 37°C prior to the addition of IPTG (0.1mM) and continued incubations for another 1.25 h. Vertical dashed lines indicate 2-fold change cutoffs of 2, while the horizontal dashed lines indicate a *P* value of 0.01. **(C and D)** Tables provide specific Log2-fold change values and brief annotations for genes highlighted in the volcano plots in A and B, respectively. Results are from 4 independent replicates; *P*<0.01 for all highlighted genes noted, as assessed by Wald tests via DESeq2.

**Table 2.** Differentially expressed genes observed in both *micC* KO and *micC* OE strains versus wild-type UTI89*.

| Gene<sup>†</sup> | KO/WT<br>(Log<sub>2</sub> fold<br>change) | OE/WT<br>(Log<sub>2</sub> fold<br>change) | Annotation |
| --- | --- | --- | --- |
| *acs* | -4.136 | -2.004 | Acetate--CoA ligase |
| *agaS* | -0.475 | -2.614 | AgaS family sugar isomerase |
| *aldA* | -2.827 | -1.197 | aldehyde dehydrogenase |
| *alsA* | -1.658 | 1.905 | D-allose ABC transporter ATP-binding<br>protein AlsA |
| *aspC* | -0.465 | 0.546 | Aspartate transaminase |
| *btuB* | -0.599 | 0.781 | TonB-dependent vitamin B12 receptor<br>BtuB |
| *cadA* | 0.811 | 1.986 | Lysine decarboxylase CadA |
| *cheA* | 0.602 | 1.881 | Chemotaxis protein CheA |
| *cheB* | 0.529 | 2.391 | Protein-glutamate methylesterase/protein<br>glutamine deamidase |
| *eamB* | 0.683 | -0.816 | Cysteine/O-acetylserine transporter |
| *fimA* | 0.950 | 2.580 | Type 1 fimbrial major subunit |
| *fimB* | -1.101 | 2.030 | Type 1 fimbria switch DNA invertase |
| *flgC* | -0.724 | 2.817 | Flagellar basal body rod protein FlgC |
| *flgD* | -0.353 | 3.981 | Flagellar hook assembly protein FlgD |
| *gabD* | -1.661 | -2.742 | NADP-dependent succinate-<br>semialdehyde dehydrogenase |
| *gabT* | -1.146 | -2.235 | 4-aminobutyrate--2-oxoglutarate<br>transaminase |
| *glcB* | -0.923 | -2.982 | Malate synthase G |
| *glcE* | -2.194 | -3.867 | Glycolate oxidase subunit GlcE |
| *glcF* | -0.899 | -2.967 | Glycolate oxidase subunit GlcF |
| *glcG* | -0.930 | -3.065 | Heme-binding protein |
| *glmS* | -1.353 | 1.250 | Glutamine--fructose-6-phosphate<br>transaminase (isomerizing) |
| *Ivy (ykfE)* | 0.656 | -1.671 | Ivy family C-type lysozyme inhibitor |
| *malF* | -3.116 | 2.832 | Maltose ABC transporter permease MalF |
| **malP<sup>†</sup>** | -0.889 | 0.841 | Maltodextrin phosphorylase |
| *manY* | -0.789 | 1.070 | PTS mannose transporter subunit IIC |
| *mdtF* | 0.451 | -2.262 | Multidrug efflux pump RND permease |
| *miaA* | -0.436 | 0.911 | tRNA (adenosine(37)-N6)-<br>dimethylallyltransferase MiaA |
| *motB* | 0.812 | 2.318 | Flagellar motor protein MotB |
| *narI* | -0.757 | -2.008 | Respiratory nitrate reductase subunit $\gamma$ |
| *nrfB* | 0.634 | 1.906 | Cytochrome c nitrite reductase<br>pentaheme subunit |
| *nrfC* | 0.533 | 1.903 | Cytochrome c nitrite reductase Fe-S<br>protein |
| *nudI* | -0.644 | 0.878 | Nucleoside triphosphatase NudI |
| **ompC<sup>†</sup>** | -0.544 | -2.140 | Porin OmpC |
| *pdeH* | 0.605 | 1.947 | Cyclic-guanylate-specific<br>phosphodiesterase |
| *pta* | -0.753 | 0.978 | Phosphate acetyltransferase |
| *rpsB* | 0.839 | 1.965 | 30S ribosomal protein S2 |
| ***sthA*<sup>†</sup>** | -1.948 | 0.794 | Si-specific NAD(P)(+) trans-hydrogenase |
| *tal* | -0.799 | 0.501 | Transaldolase |
| *tam* | -0.725 | -1.857 | Trans-aconitate 2-methyltransferase |
| *treB* | -0.677 | 3.046 | PTS trehalose transporter subunit IIBC |
| *treC* | -0.690 | 3.340 | Trehalose-6-phosphate hydrolase |
| *tsr* | 0.922 | 2.050 | Methyl-accepting chemotaxis protein |
| *tyrB* | -0.367 | 0.992 | Aromatic amino acid transaminase |
| *yagU* | -0.577 | 1.112 | Acid-induced inner membrane protein |
| *ycgZ* | -1.849 | 1.298 | Regulatory protein, may affect RcsB/C and OmpF |
| *yedE* | -1.537 | 0.759 | Selenium metabolism membrane protein YedE/FdhT |
| *yghD* | 1.147 | -1.459 | GspM family type II secretion system protein YghD |
| *yiaD* | 0.526 | 1.770 | OmpA family lipoprotein |

\*For the listed genes, $|\text{Log}_2(\text{FC})_{\text{KO}} - \text{Log}_2(\text{FC})_{\text{OE}}| \geq 1$ . FC = Fold Change; KO = UTI89Δ *micC*/pRRK1; OE = UTI89Δ *micC*/pMicC. $P < 0.05$ for KO and OE FC values relative to wild-type UTI89 carrying the empty vector pRRK1, as assessed by the Wald tests ( $n = 4$ independent replicates).
<sup>†</sup>Genes identified as potential MicC binding partners by TargetRNA3 using default settings, probability $> 0.4$ .

Several of the topmost up- and down-regulated genes in the MicC KO and OE strains are highlighted in **Figs. 7C–D**. Many of these genes are involved in the utilization of carbohydrates (including maltose, galactose, and glycerol) or are linked with motility and chemotaxis. Pathway analysis with STRING (Search Tool for the Retrieval of Interacting Genes/Proteins), which incorporates information from Gene Ontology and KEGG databases (82), confirmed and extended these observations (**Table S2**; false discovery rates < 0.05).

Transcripts that were significantly altered by at least two-fold in the *micC* deletion mutant mapped onto pathways associated with the citric acid (TCA) cycle, ABC transporters, and the metabolism of various sugars, amino acids, purines, and fatty acids. Though fewer, transcripts that were differentially regulated in the MicC OE strain were also significantly linked with multiple metabolic pathways, as well as with flagellar assembly and chemotaxis. Similar trends were revealed by STRING if we limited our analysis to the subset of genes that are differentially regulated in both the KO and OE strains relative to WT (**Table S2**, sheets labeled ‘KEGG (Shared, all)’ and ‘KEGG (Shared, abs log_2_ diff ≥ 1))’.

These RNA-Seq results suggest that MicC has additional regulatory targets beyond *ompC*. To determine if any of the altered transcripts observed in our RNA-Seq analysis are potential MicC binding partners, we employed the machine learning algorithm TargetRNA3 (83). Using the ExPEC strain EC958 as a reference, TargetRNA3 identified 81 genes with probability scores (Pr) ≥ 0.4 and *P*-values ≤ 0.05 when *micC* from UTI89 was used as a query (**Table S3**). Of these 81 genes, transcripts for 40 were significantly altered in UTI89Δ*micC*/pRRK1 relative to UTI89/pRRK1, 7 were differentially expressed in UTI89Δ*micC*/pMicC, and 4 were significant hits in transcriptional analysis of both the KO and OE strains. These 4 genes included, reassuringly, *ompC* (Pr = 0.424), as well as *malP* (encodes a maltodextrin phosphorylase involved the breakdown of maltooligosaccharides; Pr = 0.524), *aceB* (malate synthase A gene; Pr = 0.475), and *sthA* (encodes pyridine nucleotide transhydrogenase, involved in maintaining redox balance; Pr = 0.421) (84–86). Among the top 25 up- and 25 down-regulated genes in UTI89Δ*micC*/pRRK1 or UTI89Δ*micC*/pMicC (see **Table S1**), only 5 genes in addition to *ompC* were identified by TargetRNA3 as putative MicC binding partners (**Table S3**). These genes (*glpA*, *treA*, *phoH*, *lldP*, and *cadB*) each had a MicC-binding probability score on par with that calculated by TargetRNA3 for *ompC*. Transcripts for *glpA* (large subunit of the glycerol-3-phosphate dehydrogenase complex) and *treA* (periplasmic trehalase) were up 186.3- and 3.5-fold in the *micC* KO strain, respectively, while those for *phoH* (phosphate starvation-inducible ATP-binding protein) and *lldP* (lactate permease) were correspondingly downregulated 6.8 and 9.6-fold. Expression of *cadB*, which encodes a lysine:cadaverine antiporter that was previously implicated in ExPEC stress resistance and pathogenesis (87, 88), was up 5.2-fold in the *micC* OE strain. In total, these results indicate that only a subset of the many transcripts that are altered in either UTI89Δ*micC*/pRRK1 or UTI89Δ*micC*/pMicC may be direct targets of MicC.

### MicC modulates the motility of ExPEC

To phenotypically validate key findings from our RNA-Seq analysis, we focused on several of the topmost altered transcripts and pathways identified in the *micC* KO and OE strains (see **Fig. 7**, **Tables S1 and S2**). For the OE strain, these included multiple gene products involved in flagella assembly (e.g. *fliK*, *flgD*, *flgK*, *flgC*, fliD, *flgG*, *fliN*) and regulation (*flgM*, *fliA*) that were upregulated relative to the control WT strain by 2.5- to more than 17.5-fold. Relative levels of these transcripts were unaltered in the KO strain, except for the co-transcribed flagella hook assembly gene *flgD* and the basal body rod gene *flgC*, which were reduced by -1.3-and -1.7-fold, respectively. Several chemotaxis-associated transcripts (e.g. *cheA*, *cheA, cheZ, motB*, *tar*, and *tsr*) were also elevated in the MicC OE strain and, to a lesser extent, in the KO mutant. For the OE strain, these changes ranged from a 2.4-fold increase for the methyl-accepting chemotaxis gene *tar* to a 5.3-fold upturn for the associated histidine protein kinase gene *cheA*, whereas in the KO strain there was no more than a 1.9-fold increase for any of the chemotaxis-associated messages.

Following up on these observations, we tested the effects of MicC on motility by measuring the diameter of bacterial spreading over time in soft agar swim plates at 37°C. In these assays, the KO strain UTI89Δ*micC*/pRRK1 was significantly less motile than both the WT and MicC OE strains at all timepoints (**Fig. 8**). Of note, while addition of the OE construct pMicC complemented UTI89Δ*micC*, it did not render the OE strain significantly more motile than WT (**Fig. 8**). Considering previous reports indicating that the deletion of *micC* can elevate OmpC expression in K-12 *E. coli* and that OmpC levels can affect motility (89, 90), we also examined the effects of OmpC overexpression on swimming using UTI89/pOmpC. Rather than having diminished motility, as seen with the *micC* mutant, UTI89/pOmpC was notably more motile than the WT strain (**Fig. 8**). These observations indicate that the decreased motility of UTI89Δ*micC*/pRRK1 is not attributable to increased OmpC expression, in line with results showing that OmpC transcript and protein levels in UTI89 were not elevated in the absence of MicC (see **Fig. 6B** and **Table S1**). Instead, the diminished motility of UTI89Δ*micC*/pRRK1 is more likely linked with alteration of other factors, such as reduced expression of flagella-associated factors like *flgD* and *flgC* or with the dysregulation of metabolic pathways in the KO.

**Figure 8.**
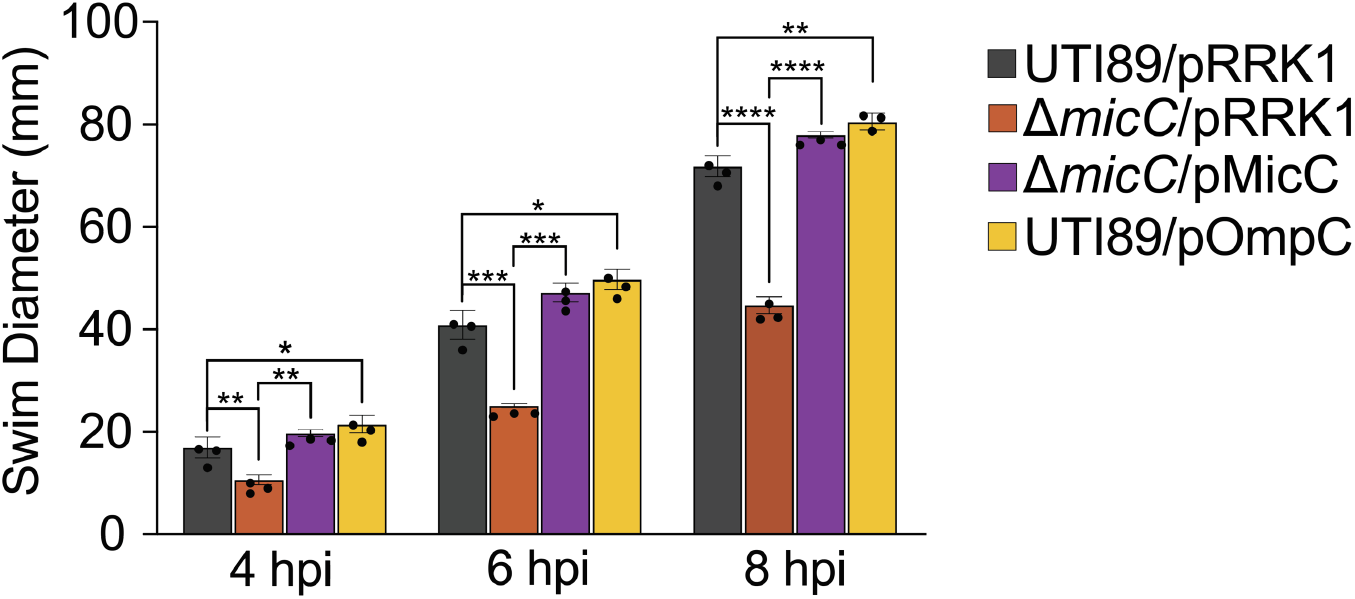
Deletion of *micC* impairs ExPEC motility. The indicated strains were inoculated into soft agar swim plates (+ 0.1 mM IPTG) and bacterial spreading was measured over the course of 8-h incubations at 37°C. Bars indicate mean diameters (± SD) of the swimming bacterial communities at 4, 6, and 8 hpi of the plates. \*\**P*<0.01, \*\*\**P*<0.001, \*\*\*\**P*<0.0001, as determined by Student’s *t* tests (n = 3 independent assays).

### MicC enhances the metabolic flexibility of ExPEC

In UTI89Δ*micC*/pRRK1, some of the most downregulated transcripts detected by RNA-Seq were associated with the utilization of maltose (*lamB*, *malE*, *malF*, *malG*, *malX*, *malY*) and galactose (*mglB*, *mglA*, *mglC*), while those involved in the catabolism of glycerol (*glpABC*, *glpD*, *glpF*, glpK, *glpT*, *glpQ*) were among the most elevated (see **Fig. 7**, **Tables S1 and S2**). Multiple transcripts linked with the TCA cycle were also repressed in UTI89Δ*micC*/pRRK1 (e.g. *acnA*, *acnB*, *fumB*, *fumC*, *gltA*, *icd*, *mdh*, *sdhCDAB*, *sucAB*, *sucCD*), though generally to a lesser extent than those associated with maltose and galactose utilization. A subset of these TCA cycle-associated transcripts (e.g. *acnA*, *sdhB*) were also downregulated in the OE strain, along with several of the genes involved in glycolate metabolism and the glyoxylate cycle (e.g. *glcDEFGB*, *aceBA*, *acs*).

While the *micC* deletion mutant grows like the wild-type strain in rich media (LB, see **Fig. 1A**), our RNA-Seq results suggest that MicC may be important under more nutrient-limiting conditions. To test this possibility, we used dilution spot assays to assess growth of the KO and WT strains on agar containing glucose, maltose, galactose, or glycerol as the sole carbon source. In these assays, UTI89Δ*micC*/pRRK1 grew significantly less robustly than UTI89/pRRK1 on all tested media (**Fig. 9**). Growth defects observed with the deletion mutant were complemented by leaky expression of MicC via pMicC. In total, these results indicate that MicC can affect multiple carbon utilization pathways, facilitating the growth of UTI89 when available nutrients are restricted.

**Figure 9.**
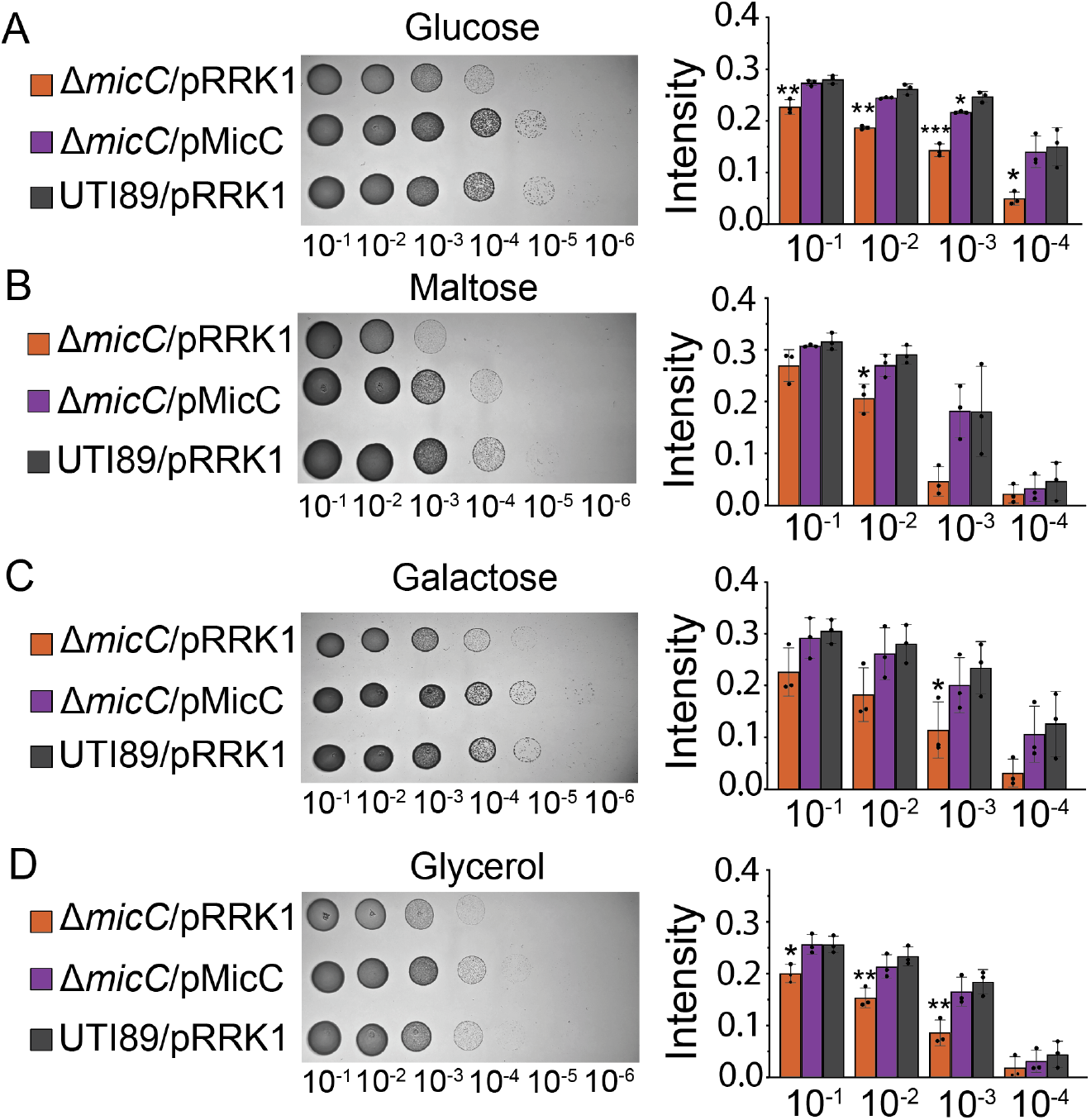
MicC facilitates growth of ExPEC on multiple single sugar carbon sources. Overnight cultures UTI89/pRRK1, UTI89Δ*micC*/pRRK1, and UTI89Δ*micC*/pMicC normalized to an OD_600_ of 0.5, serial diluted, and spotted onto M9 minimal media agar plates containing 5.55 mM of **(A)** glucose, **(B)** galactose, **(C)** glycerol, or **(D)** maltose as the sole carbon source. Plates were imaged after a 12-h incubation at 37°C **(left column)**, and spots from 3 independent replicates were quantified **(right column)**. Dilution factors are indicated below the plate images and graphs. Bars in graphs indicate mean values ± SD. \**P*<0.05; \*\**P*<0.01, \*\*\**P*<0.0005 as calculated by Welch’s *t* tests.

## DISCUSSION

ExPEC are versatile pathogens capable of adapting to diverse and rapidly changing environments. The adaptability of ExPEC and other bacteria is influenced by multiple processes, including the post-transcriptional regulation of gene expression by sRNAs (91–93). The use of sRNAs has several advantages over protein-based regulatory systems, including reduced metabolic costs and increased speed, both of which are likely important in stressful environments such as those encountered by bacterial pathogens during an infection (94, 95). In most cases, a single sRNA species can target several transcripts, thereby facilitating coordinated shifts in protein expression patterns in response to specific stressors or other environmental cues (2, 96). The results presented in this study implicate the sRNA MicC as a pivotal regulator of ExPEC fitness and virulence within distinct host niches, each with varying types of stressors. In murine models, we found that MicC promotes colonization of the bladder and intestinal tract by the reference ExPEC strain UTI89 and is essential to the virulence of this pathogen within the bloodstream. These phenotypes correspond with other data demonstrating that MicC enhances UTI89 survival within bladder epithelial cells, promotes motility, and modulates bacterial growth under nutrient-limiting conditions. A key remaining question is which MicC target(s) drive the diverse phenotypes associated with this sRNA.

In *E. coli*, OmpC is the sole validated target of MicC (37). As one of the major porins encoded by Gram-negative bacteria, OmpC is associated with maintaining membrane integrity and with the passive transport of sugars, ions, amino acids, and other small molecules (97, 98). MicC represses OmpC expression by base-pairing with sequences within the 5’ region of *ompC* transcripts, sterically inhibiting translation initiation (37). In K-12 *E. coli*, the deletion of *ompC* enables the bacteria to better avoid antibody-dependent killing via the classical complement pathway while also enhancing bacterial resistance to antibiotics (79, 97). Interestingly, ExPEC strains that are categorized as sequence type ST131, which currently rank among the leading causes of both UTI and bloodstream infections worldwide (25, 69), often downregulate OmpC and upregulate MicC relative to non-ST131 isolates (99). Reduced OmpC levels might give ST131 strains a selective advantage when faced with antibiotics while also enhancing pathogen resistance to other stressors like ROS, as seen with UTI89 in this study (see **Fig. 6C**) (79, 99). However, the deletion of *ompC* can also render *E. coli* more sensitive to bile salts, making them less fit within the intestinal tract (100). Furthermore, in a murine UTI model and in human urine *ompC* is among the most highly expressed genes in the reference ExPEC strain CFT073, suggesting that expression of this porin is important to the fitness of this pathogen within the urinary tract (101). Considering this information, it is perhaps not surprising that *ompC* is under positive selection within the ExPEC lineage (102).

In our assays, the various phenotypes observed with UTI89Δ*micC* and with the MicC overexpression strain are not entirely explainable by MicC effects on OmpC levels. For example, while the increased sensitivity of UTI89Δ*micC* to MV does mirror that observed when OmpC is overexpressed (see **Figs. 1B and 6C**), this is not the case in motility assays in which *micC* deletion and OmpC overexpression have opposite effects (see **Fig. 8**). In addition, though MicC overexpression in UTI89 strongly inhibits OmpC expression, as originally reported in K-12 *E. coli* (37), OmpC levels within UTI89Δ*micC* are not markedly different from those in wild-type UTI89 during growth in LB (see **Fig. 6A, B**). Such observations indicate that MicC can impact ExPEC fitness and virulence-related phenotypes independent of effects on OmpC expression. Our RNA-Seq results support this possibility, revealing that the deletion or overexpression of *micC* can profoundly alter the transcriptome of UTI89 (see **Fig. 7, Table S1**).

Among the transcripts that are significantly changed in the absence of *micC* are several that encode porin and porin-related factors, in addition to OmpC (-4.4-fold). These include transcripts for the maltoporin LamB (-11.2-fold), OmpW (-6.2-fold), OmpF (-3.3-fold), OmpD (-2.3-fold), OmpX (1.8-fold), the OmpC-like porin OmpN (1.6-fold), and the OmpA-like domain family lipoprotein YiaD (3.4-fold). Transcripts for OmpX and YiaD were also upregulated in the MicC overexpression strain by 2.6- and 3.4-fold, respectively. These porins have been implicated in a wide range of processes, including biofilm formation, phage sensitivity, host cell adhesion, anti-phagocytosis, nutrient uptake, and bacterial resistance to antibiotics, complement, ROS, and various other stressors (103–114). Some of these porins are co-regulated, often in an inverse fashion with one another, as a means of maintaining envelope homeostasis (80, 115–119). This could help explain how changes in MicC levels lead to the altered expression of multiple porins, which in turn could feasibly impact many of the phenotypes reported here for the *micC* deletion and overexpression strains.

Porins represent only a fraction of the hundreds of transcripts that were significantly altered in the *micC* KO and OE strains relative to the wild type. Determining which, if any, of these transcripts are direct binding targets for MicC requires further investigation. Strong candidates include the 49 genes that have *ompC*-like expression patterns, defined as being significantly altered in both the KO and OE strains and having at least a two-fold difference in expression levels between the two strain backgrounds (see **Table 2**). This candidate pool includes multiple transcripts associated with motility, chemotaxis, and various metabolic pathways, as already noted, in addition to several factors that act as mediators of bacterial colonization and stress resistance. These include genes involved in the biosynthesis of Type 1 pili (*fimA*, *fimB*), which are adhesive organelles linked with ExPEC colonization of both the urinary tract and gut (27, 65, 120), and genes that support anaerobic respiration and bacterial resistance to nitrosative stress (*narI*, *nrfB*, *nrfC, cadA*) (88, 121, 122), acid stress (*yagU*, *cadA*) (123, 124), and lysozyme (*ivy*; a.k.a. *yfkE*) (125). This list also includes the putative *ompF* regulator *ycgZ* and the tRNA modifier *miaA*, which acts as a tunable post-transcriptional modulator of multiple stress response and virulence phenotypes in ExPEC (66, 103).

Restricting our analysis of the 49 candidates in **Table 2** to those that are also predicted by TargetRNA3 to be MicC targets yielded only three transcripts: *ompC*, *malP* and *sthA*. This short-list of putative targets could reasonably be amended to include the lysine decarboxylase gene *cadA*, as this gene is co-transcribed with the computationally predicted MicC target *cadB*. Interestingly, the co-regulation of the lysine:cadaverine antiporter CadB with OmpC appears to promote *E. coli* resistance to acid stress by facilitating the transport of the CadA substrate lysine and its product cadaverine (126). In previous work we showed that cadaverine markedly enhances ExPEC resistance to nitrosative stress and promotes colonization of the urinary tract (87, 88). MicC may help coordinate translation of OmpC with cadaverine biosynthesis, while also enabling ExPEC to more effectively shift to alternate carbon sources via modulation of the maltodextrin phosphorylase MalP and the nucleotide transhydrogenase SthA. The latter enzyme drives the reoxidation of NADPH to help maintain cellular redox balance and to supply electron acceptors for catabolic metabolism (86). The coordinate dysregulation of *ompC*, *cadBA*, *malP*, and *sthA* is expected to have broad effects on multiple systems, and may explain many of the phenotypes observed with UTI89Δ*micC* and the MicC overexpression strains.

We initiated this study by searching for sRNAs that contribute to ROS resistance to help explain previous findings showing that deletion of the sRNA chaperone Hfq renders ExPEC substantially more sensitive to MV (17). In addition to MicC, our screen identified the archetypal sRNA Spf as a mediator of MV resistance (see **Fig. 1**). Spf represses numerous metabolic processes, which can often produce ROS as by-products (48). The lack of Spf may lead to higher levels of intrinsically generated ROS, compromising the ability of ExPEC to handle additional stress from exogenous ROS. Additionally, the deletion of Spf is expected to alter the levels of available Hfq binding sites, which could affect the regulation of many other sRNAs with roles in oxidative stress resistance (49). Interestingly, the only *in vivo* phenotype that we observed with UTI89Δ*spf* was in the murine intestinal tract, where the Δ*spf* mutant gained a slight competitive advantage over the wild-type strain at the 14-day time point (see **Fig. B**). This result may reflect modifications in the metabolic activities of UTI89Δ*spf* that allow the mutant to more effectively compete for nutrients within the crowded gut environment.

To date, several sRNAs have been identified that impact key fitness and virulence-associated ExPEC phenotypes (11–13, 41, 55, 92, 127–133). Like MicC, a few of these sRNAs, including RyfA and RyhB (12, 13, 130, 131), have broad effects on multiple bacterial processes, including stress resistance, motility, metabolism, host cell adhesion, intracellular survival, and colonization of host tissues. Our work with MicC builds on these studies, showing that both the deletion and overexpression of a single sRNA within ExPEC can have similarly disruptive effects, highlighting the importance of balancing sRNA levels for optimal effects. A clearer mechanistic understanding of how changing amounts of MicC can alter so many genes and pathways within ExPEC will require the definitive identification of any MicC binding partners beyond *ompC*, using results presented here as a guide and techniques like RIL-Seq (134, 135).

Similar to microRNAs in eukaryotes, bacterial sRNAs frequently act as molecular rheostats to fine-tune translation (135, 136). This means that phenotypes due to the absence or overexpression of specific sRNAs, while significant, are often relatively small. However, within the mouse bloodstream and other host niches, we found that the absence of MicC resulted in stark attenuation of ExPEC fitness and virulence. These findings indicate that MicC may be of great value as a therapeutic target. Recent work demonstrated the feasibility of using antisense oligonucleotides to disrupt bacterial sRNA activities (137). Advancing such technologies to target MicC or other critically important sRNAs within infected patients might be especially useful as a means to replace or potentiate antibiotics, enhancing their efficacy against the growing numbers of drug resistant ExPEC strains.

## MATERIALS AND METHODS

### Ethics statement

All animals used in this study were handled in accordance with protocols approved by the Institutional Animal Care and Use Committee at the University of Utah (Protocol number 10-02014), following US federal guidelines indicated by the Office of Laboratory Animal Welfare (OLAW) and described in the Guide for the Care and Use of Laboratory Animals, 8th Edition.

### Bacterial strains and plasmids

The bacterial strains and plasmids used in this study are described in **Table S4**. Knockout mutants were constructed in the reference ExPEC isolate UTI89 using the lambda Red recombination system as previously described (52, 138). The chloramphenicol resistance (Cam^R^) cassette was amplified from *Salmonella* strain TT23216 (17) or from the plasmid pKD3. Alternatively, the kanamycin resistance (Kan^R^) cassette was amplified from pKD4. Primers for amplification of the Cam^R^ or Kan^R^ cassettes were designed with overhanging ends containing ∼40 bp of homology to the 5’ and 3’ ends of target knockout sites. PCR products were then introduced by electroporation into UTI89 carrying pKM208, which encodes IPTG (isopropyl-ß-D-thiogalactopyranoside)-inducible lambda Red recombinase (138). Knockouts were verified by PCR using flanking primers specific for each target gene. Primers employed in this study are listed in **Table S5**. Plasmid constructs for expression of MicC and Spf under control of the P*tac* promoter were made using pRRK1, a modified version of pRR48 that lacks a Shine-Dalgarno sequence (139). The *micC* and *spf* genes were amplified from UTI89 by PCR, digested, and ligated into the Pst1 and Kpn1 restriction sites of pRRK1 to create pMicC and pSpf. To make pOmpC, the *ompC* coding sequence from UTI89 was cloned and ligated into pGEN-*lux*CDABE (140) so that *ompC* replaced the *lux* operon downstream of the *em7* promoter.

### Bacterial growth assays

UTI89 and its derivatives were grown from frozen stocks in 5 ml LB at 37°C overnight in loosely capped 20-by-150-mm borosilicate glass tubes at a 30° angle shaking at 225 rpm. Overnight cultures were adjusted to an OD_600_ of 1.0 and then diluted 1:100 into LB ± 1 mM methyl viologen (MV, Sigma-Aldrich). Growth curves were obtained using a Bioscreen C instrument (Growth Curves USA) with 200-μl cultures in 100-well honeycomb plates shaking at 37°C, as previously described (17). Overnight cultures of strains carrying plasmids for complementation or overexpression experiments were grown in the presence of ampicillin (Sigma-Aldrich; 100 μg/ml) to maintain the plasmids, but the antibiotic was not included in media used for the subsequent assays.

To further assess bacterial growth and survival in the presence of MV, OD_600_ of ∼0.5, diluted 1:100 into 5 ml of LB ± 1 mM MV in 20-by-150-mm borosilicate glass tubes, and grown at 37°C shaking at 225 rpm for 16 h.

Bacteria numbers were then quantified by plating serial dilutions.

### Host cell association, invasion, and intracellular persistence assays

Host cell association and gentamicin protection-based invasion and overnight intracellular persistence assays were carried out essentially as described previously using the human BEC line 5637 (HTB-9, ATCC) grown in RPMI medium supplemented with 10% fetal bovine serum (Sigma-Aldrich) (28, 66). For these assays, sub-confluent monolayers of 5637 cells in 24-well plates were infected with a multiplicity of infection (MOI) of 15 bacteria per host cell, using UTI89, UTI89Δ*spf*, and UTI89Δ*micC* from cultures grown statically for 24 h at 37°C in 20 ml modified M9 minimal media (6 g/l Na_2_HPO_4_, 3 g/l KH_2_PO_4_, 1 g/l NH_4_Cl, 0.5 g/l NaCl, 1 mM MgSO_4_, 0.1 mM CaCl_2_, 0.0025% nicotinic acid, 16.5 µg/ml thiamine in H_2_O, 0.1% glucose, and 0.2% casein amino acids). Bacterial contact with the host cells was expedited and synchronized by spinning the plates at 600 X g at room temperature for 5 min using a Beckman Allegra 6 centrifuge. Host cell association rates were determined after 1.5-h incubations at 37°C, and normalized to the total numbers of bacteria present at this time point.

Alternatively, after an initial 1.5-h incubation, infected monolayers were treated with media containing gentamicin (100 μg/ml) and the numbers of surviving (intracellular) bacteria present after additional 1-h or 17.5-h (overnight) incubations in the continued presence of gentamicin. Titers of viable bacteria recovered after the 1-h gentamicin treatments were used to calculate invasion rates relative to the numbers of host cell-associated bacteria. Overnight survival data were normalized to invasion rates.

### Outer membrane protein analysis

To examine the effects of MicC overexpression on biosynthesis of the major OMPs, overnight cultures UTI89/pMicC and UTI89/pRRK1 were diluted 1:100 into 10 ml LB containing 1 mM IPTG and then grown shaking at 225 rpm for 2.5 h at 37°C. Alternatively, to assess the effects of MV on OMP expression, wild-type UTI89 and *micC*, *spf*, and *ompC* deletion mutants were grown from overnight cultures in 10 ml LB to OD_600_≈0.5, prior to the addition of 1 mM MV for an additional 15- and 30-min. Next, 10 ml of each culture were centrifuged at 8,000 x *g* for 5 min to produce bacterial pellets that were then resuspended in 1 ml 20 mM Tris (pH 8.0) supplemented with 1 mM phenylmethylsulfonyl fluoride and protease inhibitor cocktail (Roche). Bacterial suspensions were then sonicated with a Misonix Probe Sonicater on ice for a total of 2 min, alternating 15 s on and 15 s off at a power setting of 3. Unbroken cells and large debris were pelleted at 9,000 × *g* for 2 min at 4°C and supernatants were transferred to new Eppendorf tubes prior to addition of 10% Sarkosyl (N-lauroylsarcosine) to a final concentration of 0.5% (v/v). Tubes were rocked for 5 min at room temperature and then spun at 4°C for 30 min at 21,100 × *g*. Supernatants were discarded and the outer membrane pellets were each resuspended in 500 μl of 20 mM Tris with 0.3 M NaCl (pH 7.4). Protein concentrations were determined using the BCA reagent system (Pierce), and equivalent protein amounts were resolved by SDS-PAGE using 10% acrylamide gels. Gels were stained using 0.05% (w/v) Coomassie Brilliant Blue R-250 (Bio-Rad; dissolved in 50% methanol, 10% acetic acid, 40% ddH_2_O) for 1 h at room temperature, and subsequently de-stained overnight in ddH_2_O with 5% methanol plus 7% acetic acid.

### UTI model

Seven- to eight-week-old female CBA/J mice or C3H/HeJ (Jackson Laboratory) were anesthetized using isoflurane inhalation and slowly inoculated via transurethral catheterization with 50 µl of bacteria (∼10^7^ CFU from 24-h static LB broth cultures) resuspended in PBS as previously described (17). At 3 d or 5 d post-inoculation, mice were sacrificed, and bladders and kidneys were harvested aseptically, weighed, and homogenized in 1 ml PBS containing 0.025% Triton X-100. Bacterial titers within the tissue homogenates were determined by plating serial dilutions on LB agar plates.

### Gut colonization assays

For these assays, a tagged version of UTI89 with a Kan^R^ resistance cassette inserted at the *attTn7* site (UTI89::Kan^R^) was used as the wild-type strain, so that it could be easily identified and distinguished from the *micC* and *spf* mutants which carried Cam^R^ cassettes. Previous studies demonstrated that UTI89::Kan^R^ behaves like the wild type within the murine gut (65–67). Individual cultures of UTI89::Kan^R^, UTI89Δ*spf*, and UTI89Δ*micC* were grown statically from frozen stocks for 24 h at 37°C in 250-ml flasks containing 20 ml of modified M9 medium. A total of 12 ml of culture (6 ml of each culture for competitive experiments) was centrifuged at 8,000 × *g* for 8 min. Bacterial pellets were then washed once and resuspended in 0.5 ml of phosphate-buffered saline (PBS). To inoculate the mouse gastrointestinal tract, 7- to 8-week-old female SPF BALB/c (Jackson Laboratory) were inoculated with 50 μl PBS containing 1 × 10^9^ CFU of bacteria by oral gavage. At various time points post inoculation, individual mice were briefly placed into unused takeout boxes for weighing and feces collection. Freshly deposited feces were collected from the boxes, weighed, and immediately added to 1 ml of 0.7% NaCl. The samples were homogenized shortly thereafter to break up the fecal pellets and then briefly centrifuged to pellet any insoluble debris. Supernatants were serially diluted and spread onto LB agar plates containing either chloramphenicol (20 μg/ml) or kanamycin (50 μg/ml) to select for growth of the relevant bacterial strains. Prior to gavage, fecal samples were analyzed to ensure that there were no endogenous bacteria present that were resistant to chloramphenicol or kanamycin. Mice were housed 3 to 5 per cage and given irradiated Teklad Global Soy Protein-Free Extruded chow and antibiotic-free water *ad libitum*. Competitive indices were calculated as the ratio of knockout over wild-type bacteria recovered in the feces divided by the ratio of knockout over wild-type bacteria in the inoculum, as noted previously (65).

### Sepsis model

UTI89, UTI89Δ*spf*, and UTI89Δ*micC* were grown from frozen stocks in 20 ml modified M9 medium without shaking at 37°C for 24 h, pelleted by centrifugation, and resuspended in phosphate buffered saline (PBS). Seven- to eight-week-old female Swiss Webster mice (Charles River) were briefly anesthetized by isoflurane inhalation. Mice were inoculated by intraperitoneal injection of 10^6^ bacteria as previously described (66), and survival of the mice was monitored over a 72-h period.

### Serum resistance assays

Frozen aliquots of pooled human sera, recovered from 3 healthy volunteers by using standard protocols approved by the University of Utah Institutional Review Board, were kindly provided by Dr. Andrew Weyrich. Care was taken to not freeze and thaw samples multiple times. Bacteria from overnight cultures grown with shaking at 37°C in modified M9 medium were pelleted by spinning at 8,000 × *g* for 1.5 min, washed twice with PBS, and resuspended in PBS to obtain ∼1 × 10^6^ CFU/ml. About 2 × 10^4^ CFU of each bacterial strain was mixed individually with modified M9 medium containing 20% serum, and 200-μl aliquots of each suspension were immediately placed in a 96-well microtiter plate and incubated with gentle shaking for 2.5 h at 37°C.

Plates were then placed on ice and surviving bacteria were enumerated by plating serial dilutions on LB agar. Control samples were treated with serum that was inactivated by heating for 30 min at 55°C. All results were normalized to input titers.

### RNA-seq analysis

Overnight cultures of UTI89/pRRK1, UTI89Δ*micC*/pRRK1, and UTI89Δ*micC*/pMicC were grown from frozen stocks in LB containing ampicillin (Sigma-Aldrich; 100 μg/ml) overnight at 37°C with shaking (225 rpm) and then diluted 1:100 into 5 mL LB (without any added antibiotics). Next, these cultures were grown shaking for 2 h prior to the addition of IPTG to a final concentration of 0.1 mM. After continued incubations for another 1.25 h, 1.7 ml of each culture was pelleted, and RNA was extracted using a PureLink^TM^ RNA Mini Kit (Invitrogen) and TRIzol (Life Technologies). After rRNA depletion, RNA sequencing was performed with 3 to 4 independent replicates using the Genomics Core at the University of Colorado Anschutz with an Illumina NovaSEQ 6000 instrument. Sequences were processed using BBduk v37.10 to trim adapters, and reads were mapped to UTI89 reference strain GCF_000013265 using bowtie2 v2.5.3 (141). Samtools v1.20 was used to read features (142) and differential gene expression was analyzed using the DESeq2 package in R v.4.4.0 (143).

Gene and pathway analyses were performed using STRING v12.0 (82) and EcoCyc (144). TargetRNA3 was used to identify potential MicC binding partners that matched with any significant hits obtained in the RNA-Seq analysis with either the *micC* knockout or overexpression strains. For this analysis, the ExPEC strain EC958 was used as the reference genome, as it was the most closely related strain to UTI89 available for searches with the TargetRNA3 algorithm (83). Default parameters for each program were used. Complete RNA-Seq data are available via the Gene Expression Omnibus database (https://www.ncbi.nlm.nih.gov/geo/) under accession number GSE337028.

### Motility assays

Cultures of UTI89/pRRK1, UTI89Δ*micC*/pRRK1, and UTI89Δ*micC*/pMicC were grown shaking overnight at 37°C in 5 mL LB + ampicillin (100 μg/ml) and then brought to an OD_600_ of 1.0. Next, 2 uL aliquots of each culture were inoculated just below the surface of a soft agar swim plate (0.25% agar in LB + 0.1 mM IPTG).

Swim plates were incubated at 37°C and the diameter of bacterial spreading was measured in millimeters at 4, 6, and 8 h post-inoculation.

### Bacterial dilution spot tests

UTI89/pRRK1, UTI89*micC*/pRRK1, and UTI89Δ*micC*/pMicC were grown from frozen stocks, shaking overnight at 37°C in 5 ml modified M9 medium. Cultures were then normalized to an OD_600_ of 0.5 and serial diluted prior to spotting 5 μl aliquots onto M9 minimal media agar plates containing 5.55 mM of glucose, galactose, glycerol, or maltose as the sole carbon source. Following a 12-h incubation at 37°C, plates were imaged using a Pixel 7 smartphone and areas of bacterial growth within each spot were quantified using CellProfiler (145).

### Statistical analysis

*P* values were determined by Welch’s and Student’s *t* tests, Log-rank Mantel Cox tests, or by Mann-Whitney U tests performed using Prism 8 software (GraphPad Software). Values of less than 0.05 were defined as significant.

## ACKNOWLEDGEMENTS

This work was supported in part by National Institutes of Health grants GM134331, AI135918, AI088086 and AI095647 to MAM, and by Department of Defense award W81XWH-22-1-0800 (SC210103) to WJB and MAM. Additional support was provided by University of Utah Undergraduate Research Opportunities Program scholarships to KEN and AAM, NIH Genetics T32 training grant GM007464 to OJM, and NIH Microbial Pathogenesis T32 training grant AI055434 to MGB and RRK. MGB also received support from the Federal Ministry for Research, Technology and Space (BMFTR: www.bmftr.bund.de), Germany, Project FKZ 01K12012 ‘RFIN – RNA-Biologie von Pilzinfektionen’. The funders had no role in study design, data collection and interpretation, or the decision to submit the work for publication.

