## Supplementary material for "The Small RNA MicC is a Multifunctional Regulator of Extraintestinal Pathogenic *Escherichia coli* Fitness across Multiple Host Niches": Table S4

**Table S4.** Bacterial strains and plasmids

| Strain or Plasmid | Description | Source or Reference |
|---|---|---|
| **Strains** | | |
| UTI89 | ExPEC reference strain (cystitis isolate) | (1) |
| TT23216 | *Salmonella* template strain used to amplify the $Cam^R$ cassette | (2) |
| *Recombinant Strains* | | |
| UTI89::$Kan^R$ | UTI89 with a $Kan^R$ resistance cassette inserted at the *attTn7* site | (3) |
| UTI89Δ*dsrA* | UTI89 *dsrA::*$Cam^R$ | This work |
| UTI89Δ*micC* | UTI89 *micC::*$Cam^R$ | This work |
| UTI89Δ*micF* | UTI89 *micF::*$Cam^R$ | This work |
| UTI89Δ*oxyS* | UTI89 *oxyS::*$Cam^R$ | This work |
| UTI89Δ*rhyB* | UTI89 *rhyB::*$Cam^R$ | This work |
| UTI89Δ*rprA* | UTI89 *rprA::*$Cam^R$ | This work |
| UTI89Δ*spf* | UTI89 *spf::*$Cam^R$ | This work |
| UTI89Δ*ompC* | UTI89 *ompC::*$Kan^R$ | This work |
| UT89Δ*nmpC* | UTI89 *nmpC::*$Kan^R$ | This work |
| UTI89Δ*phoE* | UTI89 *phoE::*$Kan^R$ | This work |
| UTI89Δ*spf*Δ*micC* | UTI89 *spf::*$Kan^R$*/micC::*$Cam^R$ | This work |
| **Plasmids** | | |
| pKM208 | Encoded IPTG-inducible lambda Red recombinase expression; $Amp^R$ | (4) |
| pKD3 | Template plasmid used to amplify the $Cam^R$ cassette | (5) |
| pKD4 | Template plasmid used to amplify the $Kan^R$ cassette | (5) |
| pRR48 | Contains IPTG-inducible P*tac* promoter upstream of an MCS; *$Amp^R$* | (6) |
| pRRK1 | Modified pRR48 lacking a Shine-Dalgarno sequence (used for overexpression of sRNA); $Amp^R$ | This work |
| pSpf | *Spf* from UTI89 cloned into the MCS of pRRK1; $Amp^R$ | This work |
| pMicC | MicC from UTI89 cloned into the MCS of pRRK1; $Amp^R$ | This work |
| pGEN-MCS | High-retention plasmid containing an empty multiple-cloning site; $Amp^R$ | (7) |
| pGEN-*lux*CDABE | pGEN-MCS encoding the luciferase operon under control of the *Em7* promoter; $Amp^R$ | (7) |
| pOmpC (pBF34) | Luciferase operon in pGEN-*lux*CDABE replaced with *ompC;* $Amp^R$ | This work |
