## Supplementary material for "The Small RNA MicC is a Multifunctional Regulator of Extraintestinal Pathogenic *Escherichia coli* Fitness across Multiple Host Niches": Table S5

**Table S5.** Oligonucleotides used in this study

| Primer[a] | Sequence (5'-3')[a] |
|---|---|
| *Knockout primers* | |
| DsrA-KO-F | TTCAGCGTCTCTGAAGTGAATCGTTGAATGCACAATAAAACACCAAACACCCCC CAAAACC |
| DsrA-KO-R | TATTTTCTTGTCAGCGAAAAAAATTGCGGATAAGGTGATGCACACAACCACACC ACACCAC |
| DsrA-Conf-F | GAAAGCGAAGTTCATCGCA |
| DsrA-Conf-R | ATTATCAAAGATGATTTTTTCGGG |
| MicC-KO-F | AAAATTATACTTTTAATTTTCTATACGTTATTCTGCGCGGCACCAAACACCCCCC AAAACC |
| MicC-KO-R | TTAAATGCTCTGGATAAGGATTATCCAATTCTA AAAAAAACACACAACCACACCACACCAC |
| MicC-Conf-F | GCCTTTCATCCCCATTTTG |
| MicC-Conf-R | ACTGGAAGAAACGTTACTTCACG |
| MicF-KO-F | TGTCAAAACAAAACCTTCACTCGCAACTAGAATATCTTCCCACCAAACACCCCC CAAAACC |
| MicF-KO-R | CACAGAATAATGAAAAGTGTGTAAAGAAGGGTAAAAAAACACACAACCACACC ACACCAC |
| MicF-Conf-F | TGTTTCAGAATGTAAATGAAAGGG |
| MicF-Conf-R | AGATGTACAAGCGCCATTTTG |
| OxyS-KO-F | CTATCAGGCTCTCTTGCTGTGGGCCTGTAGAATAAAAAAACACCAAACACCCCC CCAAAACC |
| OxyS-KO-R | GATTATCCCTATCAAGCATTCTGACTGATAATTGCTCACACACACAACCACACC ACACCAC |
| OxyS-Conf-F | TCCTTTGCTCCGATCGTAAC |
| OxyS-Conf-R | GGCTAACGTGGCAGGAATC |
| RhyB-KO-F | CTTCCCGAGGATAAATTGAGAACGAAAGGTCAAAAAAAAACACCAAACACCCCC CCAAAACC |
| RhyB-KO-R | GTGTTGGACAAGTGCGAATGAGAATGATTATTATTGTCTCCACACAACCACACC ACACCAC |
| RhyB-Conf-F | AGTACATACGGCAGATGGTAACG |
| RhyB-Conf-R | GTGAATCTGCCTGATGGCTT |
| RprA-KO-F | TCTGATCGACGCAAAAAGTCCGTATGCCTACTATTAGCTCCACCAAACACCCCC CAAAACC |
| RprA-KO-R | TGAGGGGCGAGGTAGCGAAGCGGAAAAATGTTAAAAAAAACACACAACCACAC CACACCAC |
| RprA-Conf-F | AAACCGAATAAGTAATTTCTCATCAG |
| RprA-Conf-R | CATCAATAGTCATGGCAAAAATAT |
| Spf-KO-F | ATGCTTTCTGAACTGAACAAAAAAGAGTAAAGTTAGTCGCCACCAAACACCCCC CAAAACC |
| Spf-KO-R | CATGGCGTATCAGGCATTACGGATCTTTTCTTTCGCCCAACACACAACCACACC ACACCAC |
| Spf-Conf-F | GAAAACTGGGATCAGGCG |
| Spf-Conf-R | TTTAGCGAGATGCAGCCTG |
| OmpC-KO-F | GCCGACTGATTAATGAGGGTTAATCAGTATGCAGTGGC CACCAAACACCCCCCAAAACC |
| OmpC-KO-R | GGGCCTGCGGGCCCTTTTTTCATTGTTTTCAGCGTACAAA CACACAACCACACCACACCAC |
| OmpC-Conf-F | GGATTATTCTGCATTTTTGGGG |
| OmpC-Conf-R | GTCGCAAGAGTACACCAAAAAA |
| PhoE-KO-F | AACTAAAACTTACATCTTGAAATAATCACATTGATTAGTGTGTAGGCTGGAGCT GCTTCG |
| PhoE-KO-R | AGGCCGGATAAGGCGTTCACGCCGCATCCGGCAATATTCACATATGAATATCC TCCTTAG |

| | |
|---|---|
| PhoE-Conf-F | CGTAGCGGCAGTTGTTTACG |
| PhoE-Conf-R | TAGAGCATCTCTTCCGGCCT |
| NmpC-KO-F | CTACTTCACAAATTAAATGAGAAACTAAAACTTACATCTTGAAATAATCACATTG ATTAGTGTGTAGGCTGGAGCTGCTTCG |
| NmpC-KO-R | GGCTCCACTTATATGTTGCGGAGGCAAAGCCTCCCGCAACATATCTTTTTCGTA AGTCAGCATATGAATATCCTCCTTAG |
| NmpC-Conf-F | TGTTCAGACAAGAGCCATGAA |
| NmpC-Conf-R | AAAAATGGGGCGTGGAAACG |
| *Cloning primers* | |
| OmpC-pGEN-F | GATACTACGTAATGAAAGTTAAAGTACTGTCCCTCCTGG |
| OmpC-pGEN-R | GATACGCTAGCTTAGAACTGGTAAACCAGGCCCAG |
| MicC-pRRK1-F | CGCGCTGCAGGTTATATGCCTTTATTGTCA |
| MicC-pRRK1-R | CGCGGGTACCGCTCTGGATAAGGATTATCCAATTC |
| Spot42-pRRK1-F | CGCGCTGCAGGTAGGGTACAGAGGT |
| Spot42-pRRK1-R | CGCGGGTACCTTCTTTCGCCCAATA |

[a] F, forward primer; R, reverse primer; KO, knockout primer; Conf, confirmation primer.
